# Characterization of a strain-specific CD-1 reference genome reveals potential inter- and intra-strain functional variability

**DOI:** 10.1101/2022.12.05.519186

**Authors:** Yoon-Hee Jung, Hsiao-Lin Wang, Samir Ali, Victor G. Corces, Isaac Kremsky

## Abstract

**Background:** CD-1 is an outbred mouse stock that is frequently used in toxicology, pharmacology, and fundamental biomedical research. Although inbred strains are typically better suited for such studies due to minimal genetic variability, outbred stocks confer practical advantages over inbred strains, such as improved breeding performance and low cost. Knowledge of the full genetic variability of CD-1 would make it more useful in toxicology, pharmacology, and fundamental biomedical research.

**Results:** We performed deep genomic DNA sequencing of CD-1 mice and used the data to identify genome-wide SNPs, indels, and germline transposable elements relative to the mm10 reference genome. We used multiple genome-wide sequencing data types and previously published CD-1 SNPs to validate our called variants. We used the called variants to construct a strain-specific CD-1 reference genome, which we show can improve mappability and reduce experimental biases from genome-wide sequencing data derived from CD-1 mice. Based on previously published ChIP-seq and ATAC-seq data, we find evidence that genetic variation between CD-1 individuals can lead to alterations in transcription factor binding. We also identified a number of variants in the coding region of genes which could have effects on splicing and translation of genes.

**Conclusions:** We have identified millions of previously unidentified CD-1 variants with the potential to confound studies involving CD-1. We used the identified variants to construct a CD-1-specific reference genome, which can improve accuracy and reduce bias when aligning genomics data derived from CD-1 individuals.

## Background

CD-1 is a commonly used outbred mouse, derived in Switzerland from two male and seven female albino mice from a non-inbred stock. CD-1 was imported into the United States in 1926, and was stably established at Charles River lab in 1959 [1]. Charles River lab uses a specific breeding program to minimize inbreeding and random genetic drift that could lead to divergence among colonies bred in different facilities worldwide.

Outbred stocks provide a good model for outbred human populations, but inbred mouse strains are typically preferred in toxicology, pharmacology, and fundamental biomedical research due to lower genetic and phenotypic variability. However, outbred stocks like CD-1 are commonly used in these settings, since they confer practical advantages over inbred strains, such as improved breeding performance and low cost. Knowledge of the full genetic variability of CD-1 would therefore make it more useful in toxicology, pharmacology, and fundamental biomedical research. [2]

A substantial amount of genetic variation exists between CD-1 individuals, and the potential to confound mapping of CD-1 genomics data has been previously reported [3]. However, the prior study only examined single nucleotide polymorphisms (SNPs) previously identified in other mouse strains, and to date, a full, genome-wide identification of genetic variants that exist between CD-1 individuals has not been done. In addition, the standard reference genome in mouse was derived from the inbred C57BL/6J strain, and the full set of genetic variants between CD-1 and C57BL/6J are not known. Mapping of transcriptomic and genomic data derived from CD-1 mice to the standard mouse reference genome is therefore subject to inaccuracies and potentially confounding effects. A major focus of this study was therefore to perform a genome-wide identification of CD-1 genetic variants, both between CD-1 individuals, and between CD-1 and C57BL/6J. For simplicity, throughout the rest of this paper, we will refer to variants between CD-1 individuals as “non-uniform”, whereas variants between CD-1 and C57BL/6J in which all CD-1 individuals sequenced have the same allele are referred to as “uniform”.

We performed deep genomic DNA (gDNA) sequencing in order to map both the uniform and non-uniform variants of CD-1, including SNPs, insertions and deletions (indels), and germline transposable elements (TEs). We then used the identified variants to construct a strain-specific CD-1 reference genome, and associated annotation files, including genes and the full set of identified CD-1 variants. We validated the CD-1 reference genome and the identified variants by comparative mapping of a wide range of CD-1 genomics datasets between the CD-1 and standard mm10 reference genomes. Mapping genomics data to the CD-1 genome rather than mm10 resulted in greater mappability and higher quality of results for all datasets examined.

We found evidence of altered transcription factor (TF) binding between CD-1 individuals at non-uniform CD-1 SNPs, highlighting the potential confounding effects that non-uniform CD-1 variants can have on functional and phenotypic studies. This study has therefore identified millions of regions in the CD-1 genome with the potential to confound studies involving CD-1. These regions are masked with “N” in the CD-1 reference genome, eliminating outright several types of potentially confounding effects at these regions. A table of these regions in standard mm10 coordinates is also provided for researchers wishing to simply map to the standard reference genome and blacklist these confounding regions. Blacklisting of these regions is recommended in the analysis of most genomic and transcriptomic data derived from CD-1 mice.

## Results

### Sequencing of the CD-1 genome and a validation dataset

We sequenced gDNA from 5 CD-1 individuals in order to determine the variants relative to the standard reference genome (mm10). The paired-end sequencing depth was at least 24-fold coverage for all 5 samples, and with all 5 samples combined, the total sequencing depth was 142-fold coverage. After trimming of adapters and low-quality bases (see Methods), all samples showed uniformly high sequencing quality across both read pairs and passed the FastQC [4] quality control measures (Additional file 1: Figures S1A-E).

We also performed Assay for Transposase-accessible chromatin followed by sequencing (ATAC-seq) on adipose tissue from 4 CD-1 individuals under two different treatments, to be used as an independent dataset to validate the results from the CD-1 gDNA data. The treatment in this case was administration of bisphenol A (BPA) in utero to F1 mice, and then samples were extracted from the F4 generation. This is the same experimental design as was done to generate whole genome bisulphite sequencing (WGBS) data which are also used in the present study for validation purposes. For the generated ATAC-seq data, adaptor and quality trimmed reads from these data were also of high quality and passed the FastQC quality control measures (Additional file 1: Figures S1F-I). Reads per kilobase per million (RPKM) values at peaks were highly correlated across replicates of each treatment (Additional file 1: Figure S1J).

### Genome-wide CD-1 variants

Using the gDNA data from 5 CD-1 individuals, we identified SNPs, indels, and germline TEs. We compared our identified SNPs to a comparable set (see Methods) derived from ~3000 previously published CD-1 SNPs [3]. 93.2% of the comparable, previously published SNPs were also called in our study prior to final filtering (Additional file 1: Figure S1K). However, for the purposes of building a CD-1 reference genome, we excluded low frequency SNPs, and also SNPs with low coverage, since more CD-1 individuals will have the major allele in that case and on average, mapping would be optimal using the major allele. We performed a similar filter on indels (see Methods).

We divided the set of filtered SNPs and indels into two main categories: uniform and non-uniform. Uniform SNPs/indels are those in which all CD-1 individuals sequenced had the same exact sequence, which was different from mm10; non-uniform SNPs/indels are those in which there is variability between CD-1 individuals. The distributions of the frequency of the most abundant alternate allele corresponded to the frequency cutoffs used for each variant class (see Methods and Additional file 1: Figure S1L). 2.4 million and 3.9 million high-confidence uniform and non-uniform SNPs were identified, respectively; we similarly identified 380,067 uniform and 742,482 non-uniform indels, 7,154 germline TEs not previously identified in CD-1 that are not contained in mm10, as well as 10,645 germline TEs in mm10 that are not in CD-1 (Table 1). After filtering, we recovered >80% of the comparable, previously published uniform SNPs and >70% of the comparable, previously published non-uniform SNPs. (Additional file 1: Figure S1M). Taken together, the validation suggests that the majority of SNPs present in CD-1 are represented in the uniform and non-uniform SNP sets we used to construct the CD-1 reference genome, while variants with low frequencies are excluded as they would result in lower mappability at such loci on average.

**Table 1.** CD-1 variants.

|  |  |
| --- | --- |
| Uniform SNPs | 2,375,189 |
| Non-uniform SNPs | 3,935,818 |
| Uniform indels | 380,067 |
| Non-uniform indels | 742,482 |
| TEs added in Cd1 | 7,154 |
| TEs lost in Cd1 | 10,645 |

### Functional consequences of genotype variation between CD-1 individuals

Non-uniform SNPs and indels in CD-1 are regions that have the potential for functional variability between CD-1 individuals. For example, a non-uniform SNP at a transcription factor (TF) binding site could result in ablated binding in individuals that deviate too far from the consensus binding motif for the TF. The result would be a TF that is bound in some CD-1 individuals but not others. Over a million non-uniform SNPs have two alleles with a roughly equal population frequency (Additional file 1: Figure S1L); these are variants with a relatively high chance that individuals in a treatment group happen to have the same nucleotide at a SNP, whereas the control individuals share a common nucleotide that differs from the treatment animals. If unrecognized, this could lead to the erroneous conclusion that altered TF binding was due to the treatment, when its actually due to a non-uniform SNP bias. Similarly, there are >120,000 non-uniform indels with two alleles with a roughly equal population frequency (Additional file 1: Figure S1L), and these could lead to similar experimental biases.

To determine whether the non-uniform SNPs can result in variations in TF binding between CD-1 individuals, we used a CTCF ChIP-seq dataset that was taken from the liver of four male and four female CD-1 individuals [5]. Upon examination, replicate 4 of the male, and replicate 4 of the female, appeared globally to be of lower quality than the other replicates (Additional file 1: Figure S2A); thus, these two replicates were removed from consideration. We then determined peaks that were ir-reproducible between the remaining replicates (Additional file 1: Figure S2B; see Methods) and took these as candidates for regions where there could be variability in Ctcf binding between individuals. The study that generated these ChIP-seq datasets identified regions of sexual dimorphism in Ctcf binding [5], and these were excluded from the set of considered ir-reproducible peaks. We reasoned that it was possible that not all sexually dimorphic regions were identified. Therefore, we decided to consider only ir-reproducible peaks from the three male individuals for a more focused, higher-confidence analysis; this left 1,461 ir-reproducible peaks (Figure 1A).

**Figure 1.**
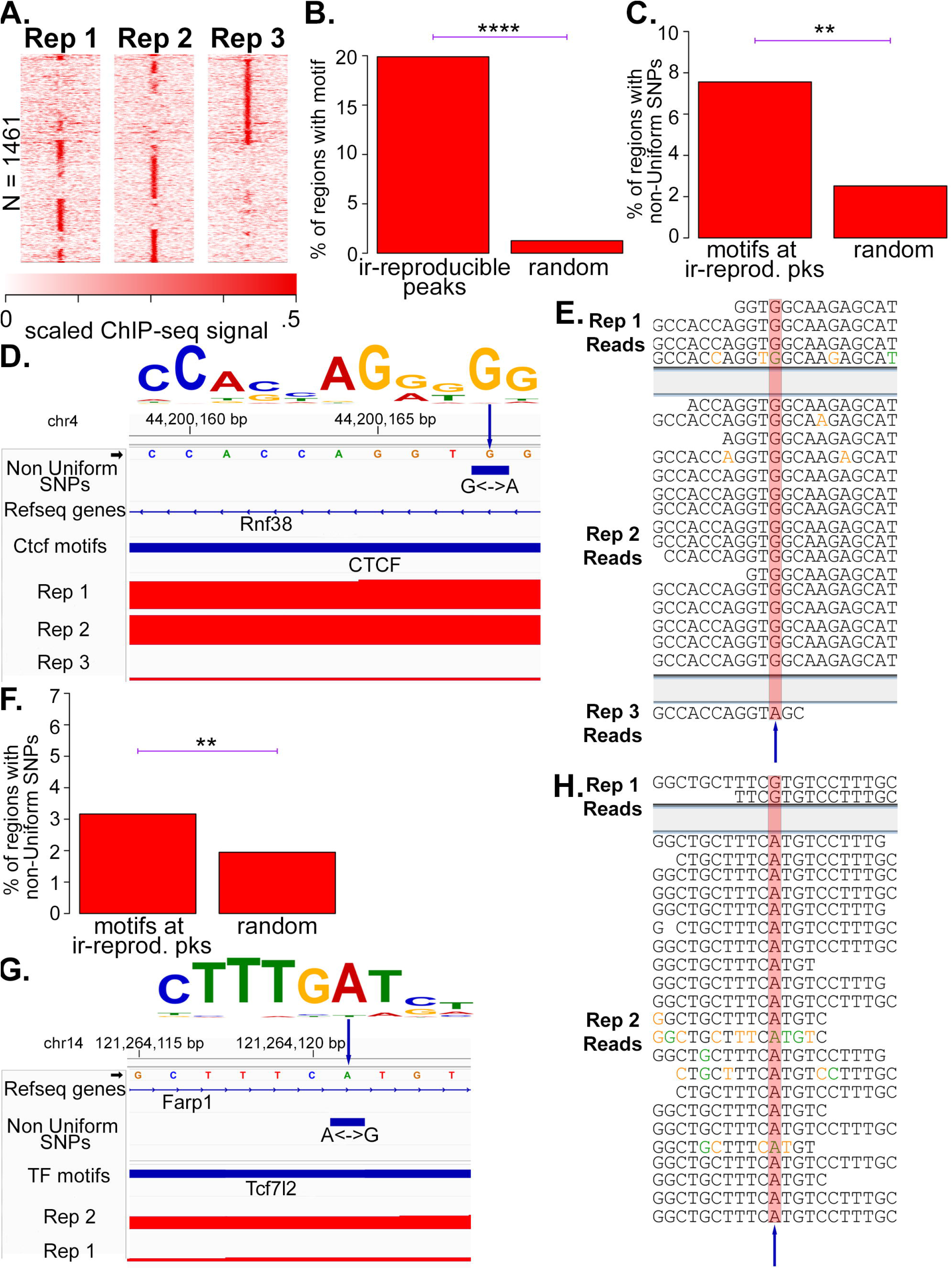
Genomic variants between CD-1 individuals with associated functional consequences. **A** Heatmaps of enrichment of Reps. 1-3 of the CTCF Male liver ChIP-seq data at ir-reproducible peaks. **B** Barplots comparing the percentage of ir-reproducible peaks (Fig. 1A) that contain a called CTCF motif to the ratio of randomly shuffled regions that contain a CTCF motif. **C** Barplots comparing the percentage of ir-reproducible peaks with a CTCF motif that overlap with a non-uniform SNP to the percentage of randomly shuffled regions that overlap a non-uniform SNP. **D** Genome browser image of Reads per million (RPM)-normalized CTCF ChIP-seq coverage around a non-uniform SNP that occurs at a highly conserved G in a called CTCF motif. The CTCF motif is pictured at top, and the arrow points to the nucleotide with the non-uniform SNP. **E** The sequenced CTCF ChIP-seq reads from the indicated replicate that map to the non-uniform SNP displayed in Fig. 1D. The non-uniform SNP is highlighted. Nucleotides are colored according to their phred sequencing quality score as follows: black for 30 or above, orange: < 30, green: < 20, blue: < 10. **F** Barplots comparing the percentage of ATAC-seq peaks with a called motif that overlap a non-uniform SNP to the percentage of randomly shuffled regions that overlap a non-uniform SNP. **G,H** Same as D,E, respectively, but for ATAC-seq data at a called Tcf7l2 motif. P-values in this figure were calculated by Fisher’s exact test, with cutoffs shown as follows: * p < .01; ** p < .001; *** p < .00001; **** p < .0000000001.

To test the hypothesis that a subset of the 1,461 ir-reproducible peaks are authentic Ctcf binding sites, we scanned for the presence of CTCF motifs at the summits of ir-reproducible peaks and compared them to a set of randomly shuffled regions. There was a highly statistically significant enrichment of CTCF motif hits at ir-reproducible peaks relative to controls (Figure 1B and Additional file 1: Figure S2C), supporting the hypothesis that a subset of the ir-reproducible CTCF peaks are authentic Ctcf binding sites. Next, to test the hypothesis that a subset of the ir-reprodcible Ctcf binding sites are ir-reproducible due to a non-uniform CD-1 SNP, we compared the proportion of regions overlapping a non-uniform SNP at motif-containing ir-reproducible peaks versus randomly shuffled controls, and indeed found a statistically significant enrichment of non-uniform SNPs near the summits of motif-containing ir-reproducible peaks (Figure 1C and Additional file 1: Figure S2D).

We next checked the specific sequences of CTCF ChIP-seq reads at ir-reproducible regions overlapping a non-uniform SNP, to see if that SNP is reflected in the ChIP-seq data in a manner that correlates with the amount of signal. Indeed, in the example shown (Figure 1D), replicates 1 and 2 have substantial ChIP-seq signal, whereas replicate 3 only has a single read. There is a non-uniform SNP at this locus, with a population-wide alternate allele frequency of 0.4, that occurs at a highly conserved G in the consensus CTCF motif. All reads from replicates 1 and 2 have a G at the SNP as expected, whereas the read from replicate 3 has an A (Figure 1E). Importantly, the A that was sequenced in replicate 3 has a phred quality score of 41, indicating the base was called at high quality and not likely to be a sequencing error.

Another example like this is shown, in which a C<->T non-uniform SNP occurs at a highly conserved C in the CTCF motif (Additional file 1: Figures S2E, F); in this example, reads from replicates 1 and 3 have a C at this position, and both have substantial ChIP-seq signal, while the single read from replicate 2 has a T at this position, with a phred score of 41. These examples corroborate the presence of non-uniform SNP’s in datasets that are independent from the data used to determine the SNPs in the first place, and establish an association between non-uniform SNPs and variability of CTCF ChIP-seq signal between CD-1 individuals. Although only a small number of reads are in the replicate where there is no peak, this is to be expected, since the ChIP-seq experiment is enriching for DNA bound to Ctcf, and so only a small number of reads from background DNA not bound to Ctcf should occur.

To determine whether this non-uniform-SNP-associated functional variability occurs more generally for other TFs, we analyzed the ATAC-seq data from adipose tissue (generated for this study) in a similar manner as was done for CTCF ChIP-seq. The majority of ATAC-seq DNA fragment lengths from these samples were less than 100 base pairs (bp) (Additional file 1: Figure S2G), which is the size-range of ATAC-seq fragments that are expected to be bound by TFs. After determining the set of TF motif-containing, ir-reproducible ATAC-seq peaks (N=521; Additional file 1: Figure S2H), we observed a statistically significant increase in the proportion of said peaks overlapping a non-uniform SNP than did randomized controls (Figure 1F). We show an example of an ir-reproducible peak with high coverage in replicate 2 but not replicate 1, with a non-uniform SNP at a highly conserved A of a Tcf7l2 motif (Figure 1G). All reads from replicate 2 contain an A at the non-uniform SNP position, whereas the two reads from replicate 1 contain a G (Figure 1H). The two G’s sequenced in replicate 1 have phred quality scores > 30, indicating that the called G is unlikely to be a sequencing error. This provides another example in which an independent dataset corroborates the presence of the called non-uniform SNP, and in addition supports the hypothesis that non-uniform SNPs can result in variable TF binding in different CD-1 individuals.

Having established evidence that the variants within CD-1 have the potential to lead to functional variability between individuals, we sought to characterize the potential, global functional consequences of both the uniform and non-uniform CD-1 variants (Figures 2A,B). The majority of both uniform and non-uniform variants occurred within introns of genes, followed by intragenic regions. The fact that more variants were identified at introns than intragenically may be due to the likely increase in mappability at introns relative to intragenic regions, since introns are typically close to mappable exons. A substantial number of uniform and non-uniform variants were identified at regions that could lead to specific functional consequences, including hundreds of uniform and non-uniform variants at splice donor and acceptor sites, and hundreds resulting in translational frameshifts (Figures 2C,D).

**Figure 2.**
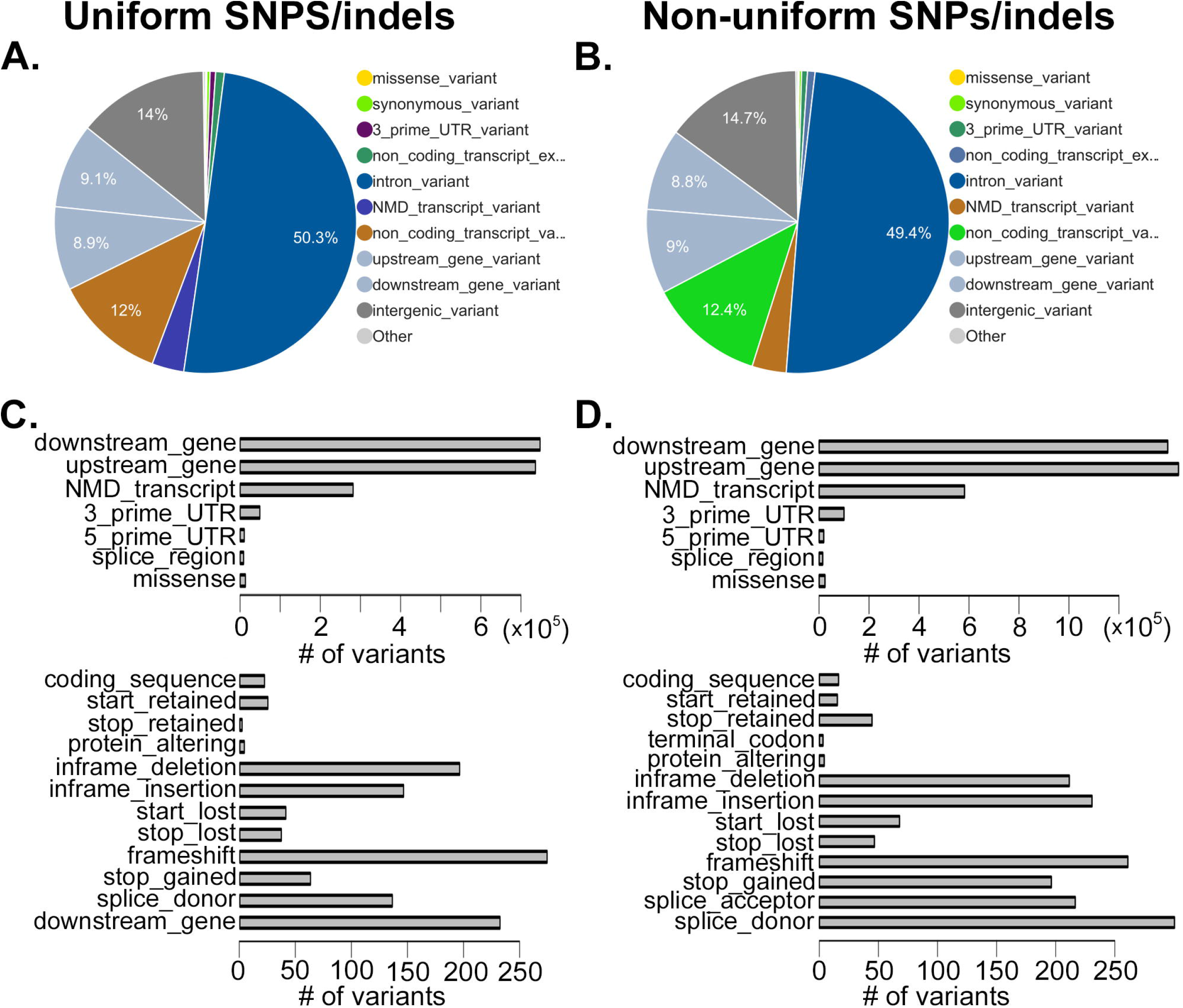
Genetic variants of the CD-1 strain. **A** Pie chart showing the distribution of the consequence types of the uniform SNPs and indels, predicted by the Variant Effect Predictor. **B** Pie chart showing the distribution of the consequence types of the non-uniform SNPs and indels, predicted by the Variant Effect Predictor. **C** Absolute counts of the indicated consequence types at uniform SNPs and indels, predicted by the Variant Effect Predictor. **D** Absolute counts of the indicated consequence types at non-uniform SNPs and indels, predicted by the Variant Effect Predictor.

### The CD-1 reference genome

Taken together, the results thus far demonstrate millions of variants with the potential to result in functional variation between CD-1 individuals and to confound nucleotide-sensitive genomics data derived from CD-1, such as ChIP-seq. The possible functional variations include but are not limited to altered TF binding, altered mRNA splicing, and translational frameshifts. Knowledge of the non-uniform CD-1 variants will be important in order to remove them from studies that use CD-1 as an experimental model, since otherwise they could confound the results and lead to functional variation that is not due to the experimental treatment. On the other hand, knowledge of the uniform CD-1 variants can enable greater accuracy when conducting sequence analyses on CD-1 mice, as there are millions of variants that separate the CD-1 genome from the standard reference genome.

We therefore created a CD-1-specific reference genome that incorporates the uniform SNPs and indels, non-uniform SNPs, and germ-line TEs identified in this study. Importantly, non-uniform SNPs are displayed as “N” in the CD-1 genome, since different CD-1 individuals have different nucleotides at these regions. This should result in more accurate alignment, since most alignment tools will not penalize mismatches at “N”, whereas, when mapping to the mm10 genome, non-uniform alleles that differ from the mm10 sequence will incur an alignment penalty.

We created chain files for converting between the CD-1 and mm10 genomes, and we used these to create a gene annotation file in CD-1 coordinates. In order to validate that the chain files are properly set up and the gene conversion is accurate, we checked the sequence at exons of the *Ctcf* and *Esrra* genes, and verified that the sequence in CD-1 coordinates matches the sequence in mm10 coordinates (Additional file 1: Figures S3A,B). In all, >99.8% of the exons annotated in the NCBI RefSeq annotation file for mm10 were successfully converted into CD-1 coordinates (Additional file 1: Figures S3C,D), and >99.7% of transcripts from mm10 are represented in the CD-1 annotation file (Additional file 1: Figures S3E,F). We have included a table of the exon IDs that were not successfully converted to the CD-1 genome, which can be used as a starting point for researchers wishing to pursue a more complete annotation of the CD-1 reference genome (Additional file 2).

### Improved mappability and accuracy of WGBS data alignment to the CD-1 genome

Given that WGBS data analysis is highly sensitive to SNPs at CpGs, we decided to align a published WGBS dataset consisting of 8 CD-1 samples to the CD-1 reference genome, both as a way to validate our called CD-1 variants, and also to explore the ability of the CD-1 reference genome to produce more accurate results when compared to aligning with the standard reference genome. Similar to the ATAC-seq data produced for this study, the WGBS samples consist of two replicates each of adipose and sperm samples of descendants of F1 mice treated in utero by BPA, as well as two replicates each of sperm and adipose samples under control conditions.

In all 8 samples, aligning to the CD-1 genome resulted in more mapped reads than when aligning to mm10 (Figures 3A,B and Additional file 1: Figure S4A). Mapping to the CD-1 genome increased the number of mapped reads by > 2% in all 8 samples, amounting to an increase of between 2 and 4 million mapped reads per sample. This occurred despite the fact that a greater number of reads of low mapping quality (mapQ < 30) occurred when aligning to mm10 (Figure 3B), demonstrating an overall improvement in both mapping quality and the number of reads mapped when mapping to the CD-1 reference.

**Figure 3.**
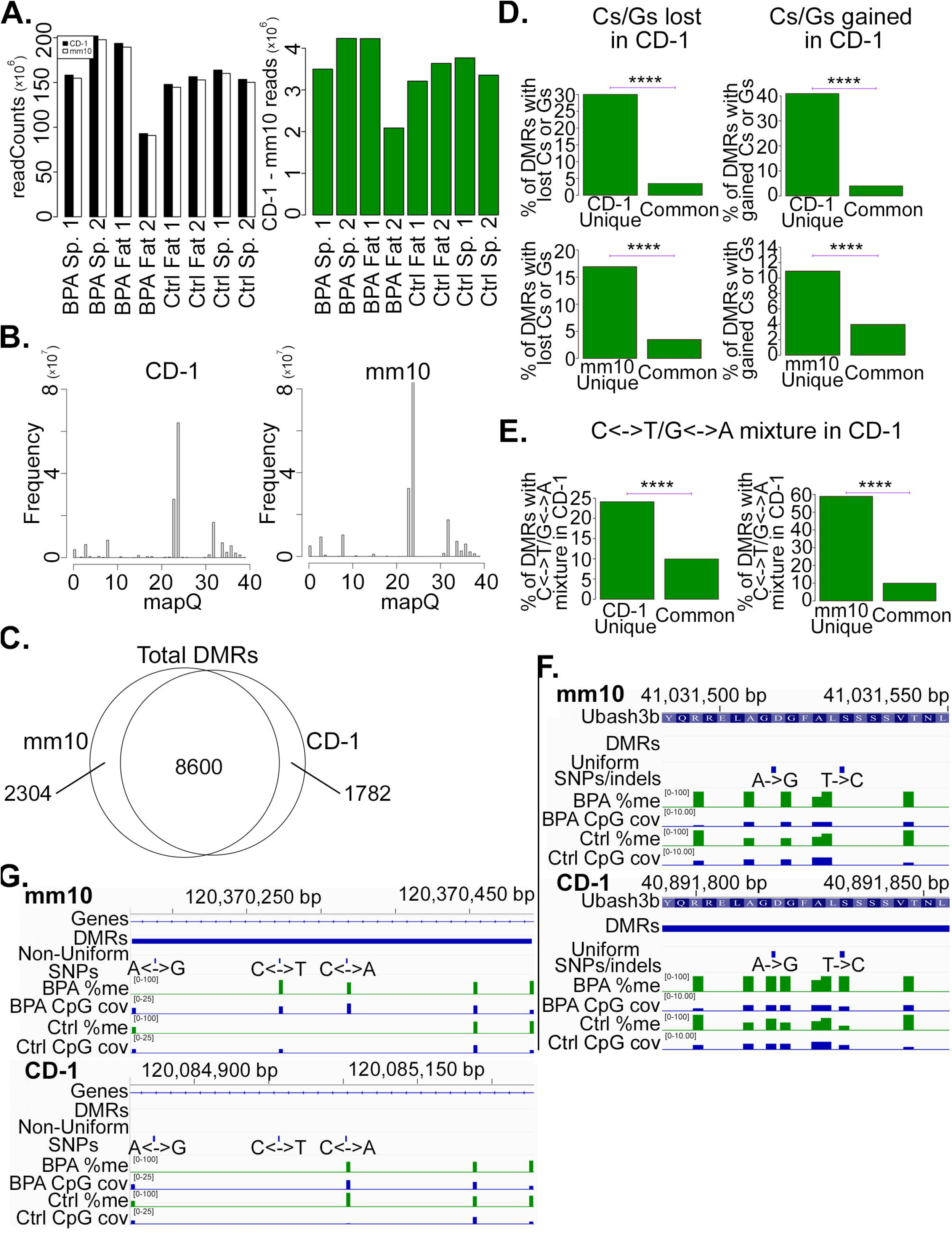
Improved mappability and accuracy of WGBS alignments to the CD-1 genome. **A** Left panel: Barplots showing the number of non-duplicate reads mapped to either CD-1 or mm10. Right panel: Barplots showing the difference between the number of nonduplicate reads mapped in CD-1 and mm10. Positive values indicate more reads mapped in CD-1 than mm10. **B** Histograms of reads mapping with mapping quality (mapQ) < 40 when mapping to CD-1 and to mm10. **C** Venn diagram showing the number of DMRs called in both mm10 and CD-1, just in CD-1, and just in mm10. All DMRs called in BPA vs. control in all samples examined in this study were combined into a single set of DMRs called after mapping to CD-1, and a single set called after mapping to mm10. **D** Barplots comparing the percentage of the indicated DMR sets overlapping with a uniform SNP in which a C or G is in mm10 but not CD-1 (top row), or comparing the indicate DMR sets overlapping with a uniform SNP in which a C or G is present in CD-1 but not mm10 (bottom row). CD-1 Unique: DMRs that are called when mapping to CD-1 but not when mapping to mm10; mm10 Unique: DMRs that are called when mapping to mm10 but not when mapping to CD-1; Common: DMRs that are called both when mapping to CD-1 and to mm10. **E** Barplots comparing the percentage of the indicated DMR sets overlapping with a non-uniform SNP of one of the indicated types. C<->T: Some CD-1 individuals have C and others T at the SNP; G<->A: Some CD-1 individuals have G and others A at the SNP. **F,G** Genome Browser views showing methylation % and WGBS read coverage at DMRs called uniquely in CD-1 and mm10, as indicated. P-values are exactly as described in Fig. 1.

Next, we called differentially methylated regions (DMRs) between the treatment (BPA) and control for each of the WGBS samples, both after aligning to mm10 as well as CD-1. The majority of DMRs were called when aligning to both mm10 and CD-1, whereas some DMRs were called only when mapping to CD-1, and still others only when mapping to mm10 (Figure 3C). 1,782 DMRs were called only when mapping to CD-1, whereas 2,304 DMRs were called only when mapping to mm10. We hypothesized that DMRs unique to one genome or the other would either occur due to the loss of false positives or to the gain of true positives when mapping to CD-1 instead of mm10. To check this, we looked at SNPs involving Cs and Gs between mm10 and CD-1, since those regions will affect the methylation calls. For example, a C in mm10 that is actually T in CD-1 would, in the absence of sequencing errors, get called as a 100% unmethylated C, potentially contributing to a false discovery. On the other hand, a T in mm10 that’s actually C jn CD-1 would get called neither as methylated nor unmethylated when mapping to mm10, but when mapping to CD-1, the region would be accurately called as either methylated or unmethylated.

There was a statistically significant enrichment of Cs and Gs that were either lost or gained in CD-1 (relative to mm10) at DMRs that were either CD-1-specific or mm10-specific (Figure 3D), supporting the hypothesis that changes in the accuracy of methylation calls are contributing to the differences in called DMRs between CD-1 and mm10. Interestingly, a higher percentage of CD-1-unique DMRs had both gained and lost Cs/Gs than did the mm10-unique DMRs, suggesting these may be contributing more to an increase in true positives than to a decrease in false positives. Upon further examination, the majority of Cs/Gs gained or lost are coming from either C->T, G->A, T->C, or A->G transitions (Additional file 1: Figure S4B), providing additional support that these are contributing to more accurate methylation calls when mapping to CD-1 than to mm10.

We also hypothesized that methylation calls based on the CD-1 reference would be more accurate than mm10 around the non-uniform CD-1 SNPs. To test this, we examined DMRs that contained a SNP that was a mixture of C and T (C<->T), or a mixture of G and A (G<->A), across different CD-1 individuals. Consistent with our hypothesis, the proportion of CD-1-unique DMRs and mm10-unique DMRs with C<->T or G<->A SNPs were both significantly higher than the corresponding proportion of DMRs that were called with both CD-1 and mm10 (Figure 3E). Nearly 60% of the mm10-unique DMRs contained at least one C<->T or G<->A SNP, suggesting that many of the mm10-unique DMRs are false positives that arise out of biases due to non-uniform C<->T or G<->A SNPs, which would be called as differentially methylated when mapping to mm10. In addition, about 25% of the CD-1-unique DMRs contain at least one C<->T or G<->A SNP. These regions do not get called as either methylated or un-methylated in CD-1, because they are masked by “N” in the CD-1 genome. Therefore, if non-uniform SNPs lie within a true DMR, they would contribute in a way that is uncorrelated with the rest of the DMR when mapping to mm10, potentially reducing the significance of the DMR and at times resulting in it not being called as significant.

We show an example of two uniform SNPs at a DMR that was called when mapping to CD-1 but not to mm10 (Figure 3F). Both of these SNPs are not recognized as CpGs in mm10, whereas in CD-1 they are, and so they have called methylation values in CD-1 but not in mm10. We also show an example of a DMR that was called when mapping to mm10 but not to CD-1 (Figure 3G). This DMR only contains 4 CpGs, one of which is at a C<->T non-uniform SNP. Since it is not clear whether T’s at this SNP were bisulfite-converted C’s, or simply individuals with T in their genome, this CpG should not get a methylation call. The unwarranted methylation call in mm10 results in a DMR getting called in mm10 amidst CpGs that do not have a major difference in methylation levels. On the other hand, in CD-1, the ambiguous reads do not contribute to DMR calling, and the potentially aberrant DMR isn’t called.

Taken together, these results show that both uniform and non-uniform SNPs can contribute in complex ways to the ultimate methylation value that is assigned at a DMR when mapping to mm10, potentially contributing to both an increase in false positives, and to a decrease in true negatives, when mapping to mm10 versus CD-1. By masking non-uniform SNPs with N’s, the CD-1 genome is protected from aberrant methylation calls that can result from biases due to non-uniform SNPs, and also increases the accuracy of methylation levels called at uniform SNPs, which are represented by incorrect nucleotides in mm10, but by the correct nucleotides in CD-1.

### Improved alignment of ATAC-seq to the CD-1 genome

While bisulfite-converted DNA is very sensitive to SNPs when used to determine DNA methylation levels, other common sequencing data types that map to the whole genome or transcriptome are more robust to SNPs. An example of such a data type is ATAC-seq, where a SNP would result in a slight mismatch score when mapping a read to the genome, but as long as the read is sufficiently long and matches the reference genome sufficiently well, it will be uniquely mapped and not influence most downstream analysis pipelines such as peak calling or transcript quantification. Therefore, the expected effect of SNPs on such sequencing data alignment is lower, but still nonzero.

To test the degree to which the CD-1 reference genome can improve the accuracy of sequence data analysis of data expected to be robust to SNPs, we used the ATAC-seq data obtained from adipose tissue of 8 CD-1 individuals described above (generated in this study). The data were mapped to both CD-1 and mm10, and peaks were called. Across all 8 samples, mapping to CD-1 resulted in a slight increase in the number of mapped reads compared to mapping to mm10 (Additional file 1: Figure S5A,B). Although the percent increase in mapped reads was small, mapping to CD-1 resulted in 50-250 thousand additional reads being mapped per sample to CD-1 than mm10 (Additional file 1: Figure S5A). The increased mappability of reads to the CD-1 genome with respect to mm10 suggests a slight improvement in mapping accuracy of the CD-1 genome even when working with sequencing data robust to SNPs.

Collectively, ~5% of all reproducible ATAC-seq peaks called were only called using either CD-1 or mm10, but not both: 2,512 peaks were called when mapping to CD-1 but not mm10, whereas 2,759 peaks were called when mapping to mm10 but not CD-1 (Figure 4A). A significantly greater proportion of peaks uniquely called either with mm10 or CD-1 contained both uniform and non-uniform variants than did peaks that were called with both CD-1 and mm10 (Figure 4B), suggesting that these variants are contributing in a substantial way to peaks that are called uniquely to either CD-1 or mm10.

**Figure 4.**
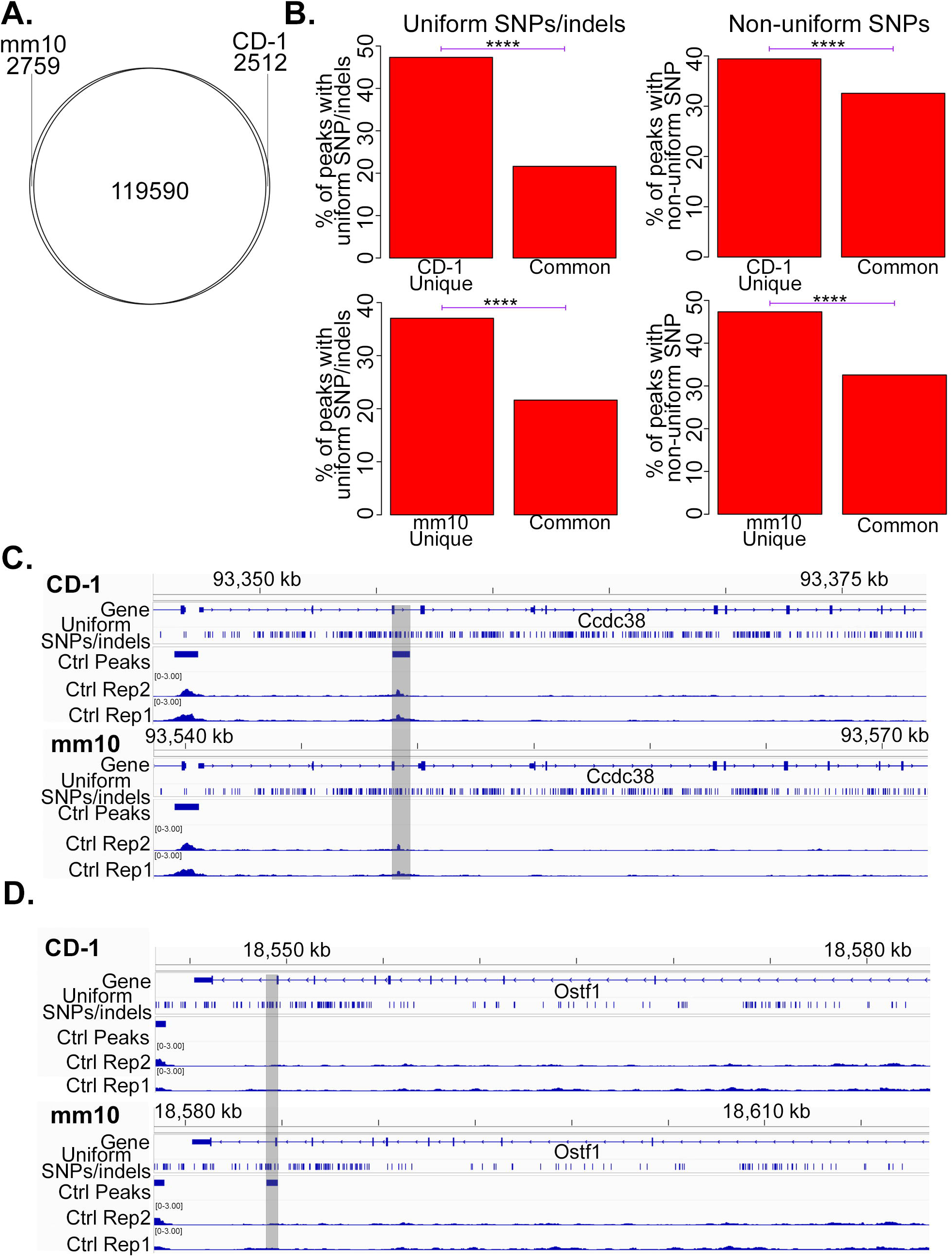
Improved accuracy of ATAC-seq alignment to the CD-1 genome. **A** Venn diagram showing the number of ATAC-seq peaks called in both mm10 and CD-1, just in CD-1, and just in mm10. Reproducible peaks (see Methods) were called separately in the BPA Adopose, Control Adipose, BPA Sperm, and Control Sperm samples, and then combined into a single pan-sample peak set for CD-1 and one for mm10. **B** Barplots comparing the percentage of the indicated ATAC-seq peak sets overlapping with uniform SNPs and indels (left column), and with non-uniform SNPs (right column). CD-1 Unique: peaks that are called when mapping to CD-1 but not when mapping to mm10; mm10 Unique: peaks that are called when mapping to mm10 but not when mapping to CD-1; Common: peaks that are called both when mapping to CD-1 and to mm10. **C** Genome browser view of an ATAC-seq peak called when mapping to CD-1 but not mm10. **D** Genome browser view of an ATAC-seq peak called when mapping to mm10 but not CD-1. In both C and D, RPM-normalized ATAC-seq coverage is displayed, and the region of interest is highlighted; the genome to which reads were mapped is indicated at the top left of each browser view. P-values are exactly as described in Fig. 1.

In an example of a peak that was called with CD-1 but not mm10 (Figure 4C), the peak region contains a number of uniform SNPs. Signal at this region can be seen to be clearly above background when examined by eye, yet was not called a peak under mm10, exemplifying an improved accuracy of peak calling when mapping to CD-1 rather than mm10. Conversely, in an example of a peak that was called with mm10 but not CD-1 (Figure 4D), examination by eye shows this region to have background-level signal. In this case, the region contains a number of non-uniform SNPs, which, when mapping to mm10, can lead some individuals to mismatch the reference genome at the SNP while other individuals match the reference genome, resulting in differential signal due to the SNP and potentially resulting in false positive calls for differential peaks.

## Discussion

Here we report the genome-wide identification of a substantial number of SNPs, indels, and TEs that vary between CD-1 and C57BL/6J, the strain used to construct the mm10 genome. In addition, we identified a substantial number of SNPs and indels that exist between CD-1 individuals, which we have demarcated with the term “non-uniform”. These non-uniform variants are particularly problematic when using CD-1 as an experimental model, because by chance, the treatment individuals could happen to have one allele at a given locus, while the control individuals have another, resulting in a bias in the number of reads mapped to this site in the treatment vs. control samples.

We identified ~3.9 million non-uniform SNPs, of which over 1 million had two alleles with a roughly equal population frequency. For biallelic variants with equal population frequencies, and assuming 2 replicates each for the treatment and control groups of an experiment, that results in a probability of ~7% that any one non-uniform SNP could have 1 allele in the 2 treatment animals, and a different allele that is common to both control animals (see Methods). Under these assumptions, the expected number of biased regions per experiment due to non-uniform SNPs would be >70,000. Similarly, we identified over 120,000 non-uniform indels with 2 alleles of roughly equal population frequencies, resulting in >8,000 expected, potentially biased regions due to non-uniform indels in each experiment. Although most non-uniform variants do not affect the coding regions of genes, 67.2% of non-uniform variants are either within introns or immediately upstream or downstream of a gene (Figure 2B), and these are regions that have the highest chance of affecting binding of TFs at promoters or enhancers and altering transcription of nearby genes. In total, we estimate that over 50,000 regions near genes will have one sequences in the two treatment animals and a different sequence that is common to the two control animals.

We used previously published ChIP-seq data to show evidence that the non-uniform SNPs can lead to variability in TF binding between individuals, providing a proof-of-principle that the non-uniform SNPs of CD-1 can lead to functional variation between CD-1 individuals. If this were to occur at one of the expected ~50,000 biased regions just described above, that would lead to a TF being bound in the two treatment animals but not in the two control animals, or vice-versa; this altered TF binding would be due to DNA sequence bias, but could be mistaken to be a result of the experimental treatment if unrecognized as a potentially biased region. 0.01% of non-uniform SNPs overlapped with an ir-reproducible peak from the adipose control ATAC-seq data from this study; assuming this to be a reasonable estimate for an average ATAC-seq experiment in most cell types, that results in at least 5 TFBS’s in any given 2X2 experiment that are expected to be bound to the two control animals but not the two treatment animals, or vice-versa, due to a non-uniform SNP, and to have a significant functional consequence. This is likely an under-estimate given the strict cutoffs used to determine the set of ir-reproducible peaks in this study. Many more non-uniform SNP loci are expected to affect only one of four replicates in an experiment, and these could also potentially confound meta analyses. Loci overlapping non-uniform CD-1 variants must therefore either be eliminated from experimental consideration, or carefully genotyped in order to draw functional conclusions from them.

In addition to functional variability, we used WGBS and ATAC-seq data to show that significant read mapping biases can arise at both uniform and non-uniform SNPs. In an effort to help eliminate non-uniform SNP biases as potential sources of erroneous experimental interpretation, we constructed a CD-1 reference genome in which the non-uniform SNPs are masked by N’s, eliminating possible biases from occurring at these loci when mapping reads to the CD-1 genome. In addition, uniform SNPs and indels between mm10 and CD-1 are replaced with the proper nucleotide sequence for CD-1 in the CD-1 reference genome. The CD-1 genome has been made available to the public, complete with gene annotations and chain files for converting genomic coordinates between CD-1 and mm10. We showed a substantial improvement in mappability and accuracy of WGBS data derived from CD-1 mice when mapping to the CD-1 reference versus mm10, and similarly showed modest improvements when mapping ATAC-seq data. More accurate DMR and peak calling is demonstrated when mapping to CD-1 rather than mm10.

When performing WGBS experiments using CD-1 mice, we recommend using the CD-1 reference genome to improve the accuracy of the results. For data types such as ATAC-seq, RNA-seq, and ChIP-seq, using mm10 should be sufficient in most cases, given the modest improvement in accuracy we observed with ATAC-seq. However, we recommend blacklisting regions that overlap with non-uniform SNPs and indels, as these represent potentially biased regions. For this purpose, we have provided a files containing all non-uniform CD-1 SNPs and indels in mm10 coordinates that can be used to simply blacklist potentially biased regions from the analysis. In cases where a treatment effect is suspected at a region overlapping a non-uniform SNP, genotyping of each individual in the experiment will be necessary to rule out SNP biases as the source of the effect. Even for data such as ATAC-seq, RNA-seq, and ChIP-seq, mapping to the CD-1 reference genome will likely confer an improved accuracy versus using mm10, as evidenced by our comparative mapping of ATAC-seq to CD-1 versus mm10. Thus, for experiments where maximizing accuracy of CD-1 sequencing data analysis is the goal, we recommend mapping to the CD-1 reference genome. Motif analysis and other sequence-based analyses will also likely benefit from improved accuracy when using the CD-1 genome for data derived from CD-1 mice.

## Conclusions

We have identified millions of previously unidentified genomic variants of CD-1 mice that have the potential to confound studies using CD-1 as a model. We constructed a reference genome specific to CD-1 that can assist researchers in obtaining more accurate results from the analysis of sequencing data derived from CD-1 mice. The results of this study can improve accuracy and reduce bias when using CD-1 in toxicology, pharmacology, and fundamental biomedical research.

## Methods

### Experimental design for genomic DNA extraction

Adult (2-5 months old) CD-1 IGS mice (Charles River Lab, USA) were used for all experiments. Mice were housed in standard cages on a 12 h light:dark cycle and given ad libitum access to food and water. Five randomly picked male animals were euthanized by cervical dislocation before the cortex were retrieved for DNA extraction and genomic library preparations. All animal experiments were approved by the Emory University Institutional Animal Care and Use Committee (IACUC).

### Genomic DNA extraction and library preparations

Dounce homogenizers were used for each cortex sample in a general lysis buffer containing 0.1 M Tris-pH8, 0.2M NaCl, 5 mM EDTA, 10% SDS, 150 mM DTT and proteinase K. The genomic DNA was extracted from the lysate using phenol:chloroform:isoamyl alcohol extraction followed by EtOH precipitation. 2 μg of genomic DNA was sheared with Diagenode Bioruptor to yield DNA fragments that are approximately 300 bp. DNA fragments were end repaired, A-tailed, and ligated to Illumina adaptors followed by PCR amplification for 7 cycles. The final libraries were size selected using AMPure XP beads, so the fragments were between the size of 200-400 bp long. Each library was subject to paired-end sequencing for 150 bp reads using Illumina NovaSeq platform.

### Experimental design for ATAC-seq data generation

Mice were maintained and handled in accordance with the Institutional Animal Care and Use policies at Emory University. All experiments were conducted according to the animal research guidelines from NIH and all protocols for animal usage were reviewed and approved by the Institutional Animal Care and Use Committee (IACUC). Mice were housed in standard cages on a 12:12 h light:dark cycle and given ad lib access to food and water. Healthy 8-week old CD-1 mice (Charles River Labs) not involved in previous procedures were used for all experiments. Gestating females (F0) were administered daily intraperitoneal injections of Bisphenol A (Sigma 239658, 50 mg/kg) or sesame oil (Sigma S3547) to prevent irritation at the injection site. Injections were performed from embryonic day 7.5 through 13.5. Offspring from different BPA-treated F0 mice were mated (no sibling breeding was used to avoid inbreeding artifacts) to produce offspring of BPA-treated mice that were not directly treated by BPA themselves, and this was done similarly for control mice. Direct, untreated descendants of BPA-treated and control-treated F0 mice were generated separately in this manner through the F4 generation, which is where samples were taken for sequencing.

### Sample extraction and library preparation for ATAC-seq

ATAC-seq was carried out using the Omni-ATAC protocol [6]. Visceral adipose tissue (1g) was placed in cold 1× homogenization buffer (320 mM sucrose, 0.1 mM EDTA, 0.1% NP40, 5 mM CaCl2, 3 mM Mg(Ac)2, 10 mM Tris pH 7.8, 1× protease inhibitors (Roche, cOmplete), and 167 μM ß-mercaptoethanol, in water), homogenized with Dounce homogenizers, and residual debris was precleared by using 80 um nylon mesh filter. Nuclei were then collected by layering with iodixanol mixture. 50,000 counted nuclei were transferred to a tube containing the transposition mix (25 μl 2× TD buffer, 2.5 μl transposase (100 nM final), 16.5 μl PBS, 0.5 μl 1% digitonin, 0.5 μl 10% Tween-20, 5 μl water) and mixed by pipetting up and down six times. Transposition reactions were incubated at 37 °C for 30 min in a thermomixer with shaking at 1,000 r.p.m. Reactions were cleaned up with Zymo DNA Clean and Concentrator 5 columns. Library amplification was done with 2x KAPA HiFi mix (Kapa Biosystems) and 1.25 μM indexed primers using the following PCR conditions: 72°C for 5 min; 98°C for 30 s; and 10-11 cycles at 98°C for 10 s, 63°C for 30 s, and 72°C for 1 min.

### Pre-processing of CD-1 genomic DNA data

Adaptors were trimmed from raw sequencing reads using trimmomatic version 0.38 [7]. The script used for this has been provided on github. All subsequent analyses using these data started from the trimmed fastq files that resulted from this step.

### Identification of germline transposable elements different between CD-1 and mm10

We first constructed a fasta file of the consensus sequence for each mus musculus TE annotated on the Dfam website [8]. We then used TEMP2 [9] insertion (version 0.1.1) to detect TE insertions in each of the 5 samples separately, using the trimmed fastq files for each sample, and the fasta file of consensus motifs as inputs. From the output of TEMP2, we made a custom script to extract all candidate germline TEs by requiring that the allele frequency be at least .495, since a germline TE must be fully present on at least one of the two alleles, and .495 rounds up to the 50% allele frequency required. In addition, candidate germline TEs were required to have at least two reads supporting the insertion. Then, we merged the trimmed fastq files from all 5 CD-1 mice into a single fastq file for each read pair, and the two merged fastq files were then input into TEMP2 insertion in order to get population-wide allele frequencies for each TE identified. We filtered TEs from the pooled analysis, and kept only those with a minimum population allele frequency of just under 10% (corresponding to the germline in 1 of the 10 alleles, allowing for a small amount of sequencing error). Finally, the filtered TEs from the population-pooled analysis were only selected if they were also on the candidate germline TE list mentioned above, ensuring that the TE is contained on at least one allele in at least one individual, with tolerance for a small amount of sequencing error. We were not very strict with our cutoffs for TEs, because we reasoned that inserting a TE that wasn’t actually a germline TE would not have much of an effect when aligning reads to a reference genome with these TEs inserted. On the other hand, with stricter cutoffs that would increase the odds of filtering out authentic germline TEs, and not having those in the genome would make it less accurate.

We also ran TEMP2 absence, specifying an insert size of 200 bp on the DNA fragments (-f 200). For the final list of TEs absent from CD-1, we required at least 10 reads that support its absence, and an estimated population frequency of 100% with the absence. After obtaining the coordinates for the final list of germline TE insertions and TEs in mm10 that are absent in CD-1, we modified the reference genome to insert the consensus sequence for each germline insertion, and removed the TE sequences for the mm10 TEs that are absent from the germline in CD-1, using RSVSim [10] version 1.26.0. Newly inserted germline TEs were first inserted into mm10 with the function simulateSV(), with the parameters sizeDups=0, bpFlankSize=0, maxIndelSize=0, maxDups=0, random=F, seed=2, percCopiedIns=1. These input parameters ensured that the consensus sequence for only the inserted TEs identified above were inserted into mm10 exactly once, at precisely the position where the insertion was identified, and no random insertions were made. We then took the fasta output from simulateSV(), and used that as the input for a 2^nd^ call of the function simulateSV() in order to remove the TEs identified as being absent from CD-1, as described above. In this case, we again used the input parameters random=F, seed=2, percCopiedIns=1 in order to ensure that only the TEs identified as absent were removed from the reference, and in precisely the position at which they were identified to have been missing from.

The resulting genome was soft masked using RepeatMasker [11] version 4.1.2-p1 with the flags “-xsmall -species mouse”. This marks repetitive regions with lower case letters for greater efficiency of analysis with a number of bioinformatics tools. The result of this was a fasta file which was the mm10 reference containing the consensus sequence for the newly identified CD-1 TEs, and with absent TEs removed; this fasta, which we will call “mm10-CD-1 hybrid” for brevity, was then used for SNP and indel calling as described below.

### SNP and indel calling

We then took the mm10-CD-1 hybrid fasta reference described above, and used GATK [12] version 4.2.0.0 to call SNPs and indels on it, based on the gDNA data, and then incorporate the called variants into the mm10-CD-1 hybrid reference genome. In this way, the deviations from the consensus sequences of the newly inserted TEs could be detected and corrected, as could SNPs and indels relative to mm10 at non-TE regions.

In order to use the gDNA data in GATK, we first mapped the trimmed fastq files for each sample to the mm10-CD-1 hybrid reference fasta. Alignments were done using bwa [13] mem version 0.7.17-r1198-dirty. PCR duplicate reads were removed from the resulting .sam files using samtools [14] markdup version 1.10. Files were converted to sorted bam format using samtools and saved for downstream analysis.

The sorted, duplicate-removed .bam files for each of the 5 CD-1 individuals were then used as inputs for a single call of GATK HaployTypeCaller, along with the mm10-CD-1 hybrid fasta file, in order to call SNPs and indels between the mm10-CD-1 hybrid genome and CD-1. Called SNPs and indels were saved into two separate files using GATK SelectVariants. SNPs were then filtered with GATK VariantFiltration with the flags “-filter “QD < 2.0” -filter “QUAL < 30.0” - filter “SOR > 3.0” -filter “FS > 60.0” -filter “MQ < 40.0” -filter “MQRankSum < −12.5” -filter “ReadPosRankSum < −8.0”” to ensure only high-confidence SNP calls. Indels were similarly filtered with the flags “-filter “QD < 2.0” -filter “QUAL < 30.0” -filter “FS > 200.0” -filter “ReadPosRankSum < −20.0”” in order to ensure high-confidence indels.

After filtering, “uniform” variants were selected using GATK SelectVariants with the flags “-select “AF > 0.8 && AC > 5” --restrict-alleles-to BIALLELIC”. This ensures that all SNPs and indels identified as uniform have an allele frequency > 80% and that more than half of the 10 total alleles are present. Uniform indels were further filtered using GATK RemoveNearbyIndels with the flag “--min-indel-spacing 1”. The final set of uniform SNPs and indels were merged into a single VCF file using GATK MergeVcfs.

“Non-uniform” SNPs were taken as the subset of filtered SNPs which were either biallelic with an allele frequency between 20% and 80% (GATK SelectVariants with the flags “-select “AF <= 0.8 && AF > 0.2 && AC > 5” --restrict-alleles-to BIALLELIC”), or multiallelic with the most frequent allele being > 20%; this was achieved using GATK SelectVariants with the flags “-select ‘AN > 5 && vc.hasGenotypes() && vc.getCalledChrCount(vc.getAltAlleleWithHighestAlleleCount())/(1.0*vc.getCalledChrCount()) > 0.2’ --restrict-alleles-to MULTIALLELIC”. Non-uniform biallelic and multiallelic SNPs were merged into a single VCF file with GATK MergeVcfs.

### SNP validation

We obtained a previously published set of CD-1 SNPs genotyped in 245 CD-1 mice [3]. From that list, we obtained a set of uniform SNPs by taking only the SNPs with a minor allele frequency (MAF) of 0 and in which the minor allele nucleotide was the nucleotide in the mm10 reference genome. Non-uniform SNPs were taken as those with MAF>0.2, and in which the nucleotide of either the minor allele or the major allele was the nucleotide present in the mm10 reference genome. The genomic coordinates of the published SNPs were done in mm8, so liftOver [15] was used to convert them to mm10, for direct comparison with our set of uniform and non-uniform SNPs.in mm10 coordinates. The uniform and non-uniform sets obtained from the previously published SNPs were combined for comparison with our full set of SNPs prior to removal of those with low allele frequencies and low coverage.

### CTCF ChIP-seq data analysis

Raw fastq files of CTCF ChIP-seq were downloaded from the NCBI GEO [16] website using their fasterq-dump tool. Fastq files were then trimmed using trim_galore [17] version 0.6.6., a wrapper for cutadapt [18] (version 2.8). Trimmed fastq files were then mapped to the mm10 reference genome using bowtie2 [19] version 2.3.5.1, and peaks were then called with macs [20] version 2.2.7.1. Peaks from all replicates were merged using bedtools [21] merge (version 2.27.1). Reads per kilobase per million (RPKM) values were then calculated for all replicates at the set of merged peaks. All peaks where the minimum RPKM across all replicates considered was < 0.5, and where the maximum RPKM across all replicates considered was >2, were selected as the set of ir-reproducible peaks. We then used bedtools shuffle to create a random set of control regions for comparison with the ir-reproducible peaks. RepeatMadker regions from mm10, as obtained from the UCSC table browser [22], were excluded from use in bedtools shuffle, with the flag “-f .5”, to ensure that no randomly shuffled region overlaps by more than 50% with a repetitive region.

Next, the mm10 sequences at all ir-reproducible and randomly shuffled peaks were extracted using bedtools getfasta, and then TF motif scanning was done at these sequences using fimo [23] version 5.3.0, using a combination of the jolma2013.meme, JASPAR_CORE_2014_vertebrates.meme, and uniprobe_mouse.meme TF motif data files provided by fimo for motif scanning. We then compared the proportion of ir-reproducible peaks with a CTCF motif hit (either human or mouse) to the proportion of random peaks with such a motif hit, using Fisher’s exact test to calculate the P-values, to perform the analysis presented in Figure 1B. We similarly compared the proportion of ir-reproducible peaks with a motif that overlap a non-uniform SNP to the proportion of randomly shuffled regions that overlap a non-uniform SNP, to perform the analysis presented in Figure 1C. P-value calculations were made using the R function fisher.test() in R version 4.0.3.

ChIP-seq read sequence content was compiled as follows. .bam files of duplicate-removed CTCF ChIP-seq alignments to mm10 were input into bamView [24] version 18.2.0, with “Colour by” set to “Base Quality”. The CTCF and reverse-compliment CTCF motif images used in the main and supplemental figures were generated using meme2images, a tool from the MEME Suite [25] version 5.3.0.

### Classification of CD-1 variants

Uniform SNPs/indels, and non-uniform SNPs/indels, were input separately into the command-line version of the Ensembl variant effect predictor [26] version 104. The gtf file, “Mus_musculus.GRCm38.102.gtf.gz” was downloaded from the Ensembl website [27] and processed according to the vep User’s Manual, for use with the –gtf flag of vep. We used the pie charts under the heading “Consequences (all)” for display in Figure 2. In addition, we extracted.the table of counts of each consequence type under “Consequences (all)” for plotting in barplots.

### Construction of the CD-1 reference genome and associated annotation files

GATK FastaAlternateReferenceMaker was used to convert to CD-1 nucleotides at the uniform SNPs and indels in the mm10-CD-1 hybrid genome. The non-uniform SNPs were used within the flag “--snp-mask”, resulting in non-uniform SNPs being masked by “N” in the final CD-1 genome. The resulting genome was then soft-masked for repeats with RepeatMasker [11], with the flags “-xsmall -species “Mus musculus”.

To facilitate conversion between the coordinate systems of mm10, the mm10-CD-1 hybrid, and CD-1, .chain files were constructed using the commands recommended by the genomewiki site [28], which makes use of several tools from the UCSC genome browser [15]: faSplit, liftUp, axtChain, chainNet, faToTwoBit, chainMergeSort, twoBitInfo, and netChainSubset, as well as blat [29], and samtools faidx, The script we used to make the .chain files is available on our Github repository page. The chain files for converting between CD-1 and mm10 coordinates have been made publicly available.

VCF files of uniform SNPs/indels, and non-uniform SNPs/indels, in mm10 coordinates were obtained via UCSC’s liftOver tool [15], by using the .chain file attained as described above to convert from the mm10-CD-1 hybrid coordinates to mm10 coordinates. These VCF files are publicly available.

An annotation .gtf file of genes was created in CD-1 coordinates as follows: a refFlat gtf file in mm10 coordinates was downloaded from the UCSC Table Browser [22], and the tool crossMap [30] was used in gff mode to convert it to CD-1 coordinates. This resulted in a simple genome annotation that was used for viewing in the Integrative Genomics Viewer (IGV) [31]. However, we observed some instances of exons that were repeated in regions of the genome that were far from the rest of the gene and most likely errenous. Therefore, in order to construct a more accurate gene annotation, we used the liftoff [32] software (version 1.6.3), which was specifically designed for accurate mapping of gene annotation files. The file “mm10.ncbiRefSeq.gtf “ was downloaded from the UCSC Genome Browser website [15] and then converted to CD-1 coordinates using liftoff with the flags, “-infer_genes -exclude_partial - polish -chroms”. We verified that the IGV screenshots in this paper, which used the simpler refFlat annotation, were also correctly annotated with the file created with liftoff. The annotation file created with liftoff has been made publicly available.

### ATAC-seq data analysis

Raw fastq files were trimmed using trim_galore as described above for CTCF ChIP-seq. Trimmed fastq files were then mapped separately to both the mm10 and CD-1 genomes as follows. Reads were mapped using bowtie2 [33] version 2.3.5.1, with the flag “-X 2000” to allow for DNA fragments up to 2 kb. Reads that did not map uniquely to one location were removed, and then reads were sorted, and PCR duplicates were removed with samtools markdup. Peaks were then called on each individual sample with macs2 [34] callpeak version 2.2.7.1. We kept only peaks that overlapped by at least 50% in the two replicates of each sample, and kept these as the set of reproducible peaks. Reproducible peaks from all samples were then merged into a single peak set separately for mm10 and CD-1, using bedtools merge.

Ir-reproducible ATAC-seq peaks were obtained as follows. Peaks from both Control Adipose replicates were merged using bedtools [21] merge (version 2.27.1). Reads per kilobase per million (RPKM) values were then calculated for each replicate at the set of merged peaks. Peaks with RPKM<1 in one replicate and RPKM>2 in the other were selected as the set of ir-reproducible peaks. The generation of randomly shuffled control regions, TF motif analysis, the analysis of SNPs at ir-reproducible peaks vs. control peaks, and the display of sequence content of the ATAC-seq reads, were all done in the same manner as described above for the CTCF ChIP-seq data.

### WGBS data analysis

Raw fastq files were trimmed using trim_galore version 0.6.6. Trimmed fastq files were then mapped separately to both the mm10 and CD-1 genomes as follows. Reads were mapped using bismark [35] version 0.22.3. PCR duplicates were then removed with the command deduplicate_bismark, and then bismark_methylation_extractor was used to extract methylation counts at CpGs. Then, for each sample, DMRfinder [36] version 0.3 was used to call DMRs of BPA vs. control for the available tissue samples.

### Estimation of the probability of experimental bias due to non-uniform CD-1 variants

In an experiment with 2 treatment and 2 control replicates, at a variant with 2 alleles, there are 2^8^=256 possible genotypes. Denoting one allele as “a”, and the other as “b”, and the genotype of the two treatments first, followed by the genotype of the two control replicates, there are 18 possible combinations where the two treatment replicates have one genotype and the two control replicates have a common genotype which differs from the treatment mice:

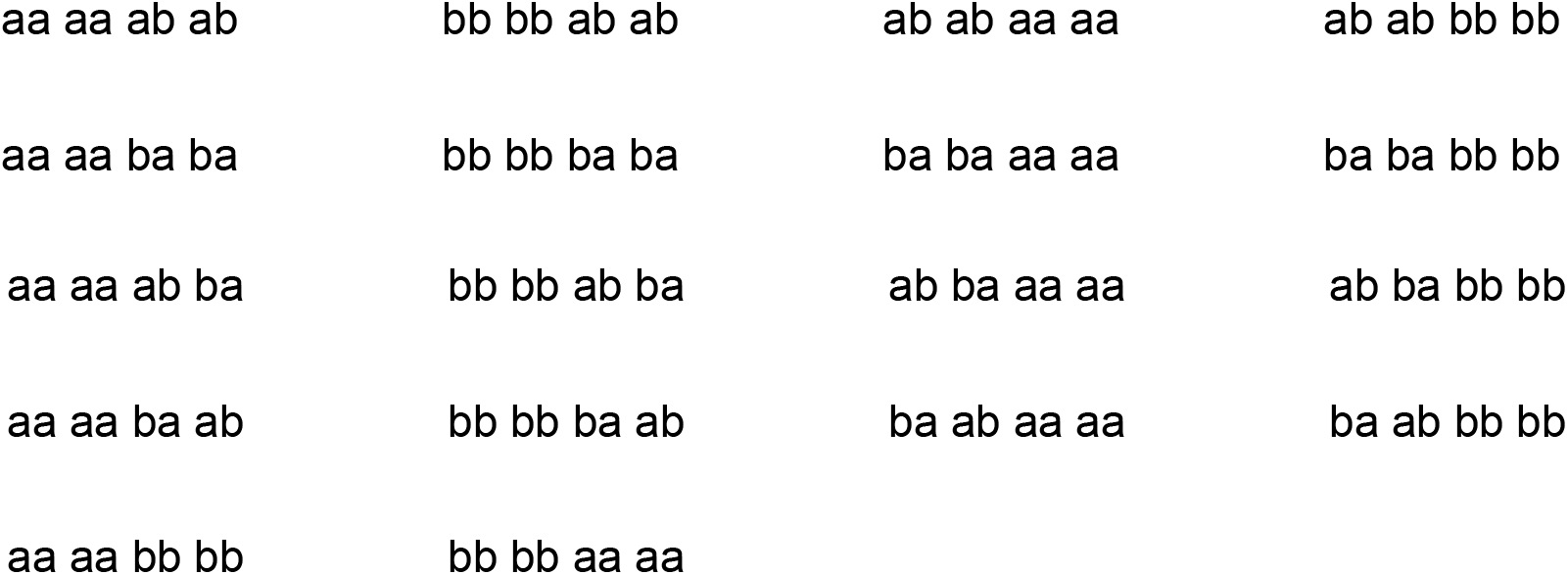

So the probability of the two treatment genotypes being the same, and differing from a common control genotype, are 18/256 = 7.03125%. The expected number of such occurrences genome wide is then just the total number of non-uniform variants times 7.03125%. Considering only non-uniform variants that are bi-allelic and where each allele has a roughly equal frequency (between 40% and 60%), this results in an expected 10^6^X7.03125%=70,312 non-uniform SNPs and 120,000X7.03125% = 8,437 non-uniform indels. To get the final estimate of 50,000, we added these two numbers together (70312+8437) and then multiplied by the percentage of non-uniform variants that were either at introns, or immediately upstream or downstream of a gene (Figure 2B).

## Declarations

### Ethics approval and consent to participate

Not applicable.

### Consent for publication

Not applicable.

### Availability of data and materials

Genomic DNA sequencing and ATAC-seq data that support the findings of this study have been deposited in NCBI GEO with the accession code GSE209816 (token: **unszskikljozril)**. All previously published data analyzed in this study are available at the NCBI GEO repository at the following locations: CTCF ChIP-seq, GSE102997 and GSE130908 [5]; WGBS, GSE149302. All custom scripts used to process and analyze the data and to obtain the results described in this paper have been deposited in the GitHub repository (https://github.com/ikremsky/Scripts-for-Characterization-of-a-strain-specific-CD-1-reference-genome...-).

### Authors’ contributions

IK and VC conceived and designed the study. YJ and HW performed experiments. IK and SA performed Bioinformatics analysis. IK, YJ, and HW wrote the manuscript. All authors approved the manuscript.

## Acknowledgements

None.

## Supplementary information

**Additional file 1: Figure S1.** Validation of sequencing data generated in this study. **Figure S2.** Related to Figure 1. **Figure S3.** Validation of the newly generated CD-q genome annotation. **Figure S4.** Related to Figure 3. **Figure S5.** Related to Figure 4.

**Additional file 2.** Ids of exons that have not yet been annotated in the CD-1 reference genome.

