## Additional file 1 for "Characterization of a strain-specific CD-1 reference genome reveals potential inter- and intra-strain functional variability"

**Figure S1**

A.
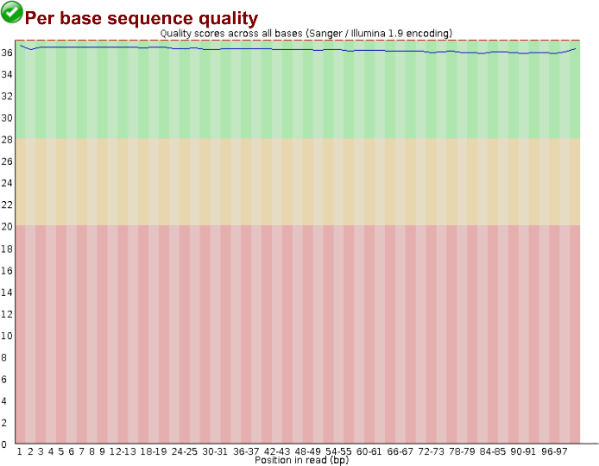

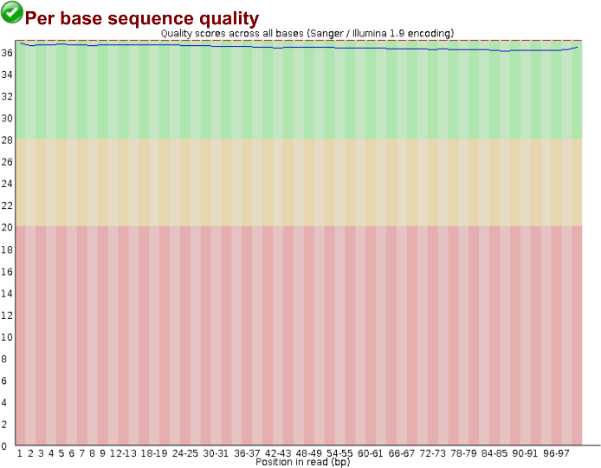
 **CD1-brain-M-2M-DNA-4**
B. **CD1-brain-M-3M-DNA-1**

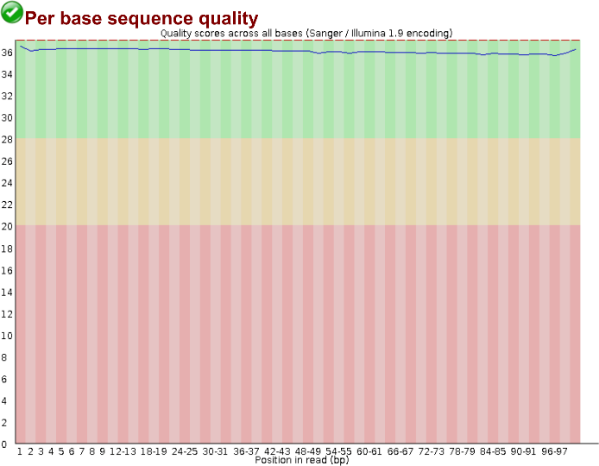

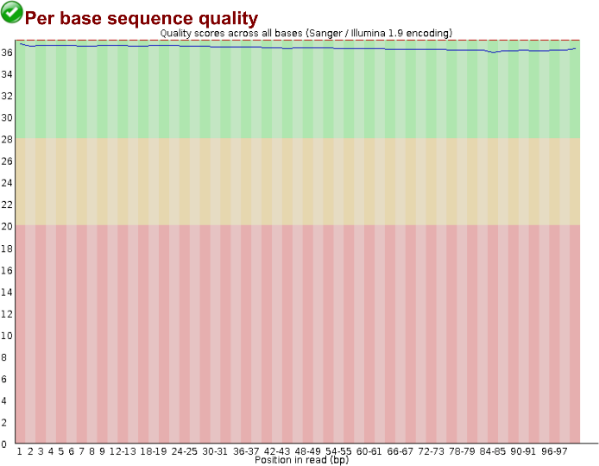

C. **CD1-brain-M-4M-DNA-2**

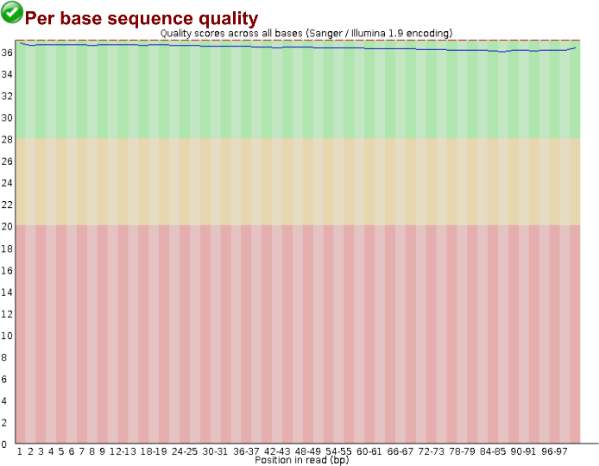

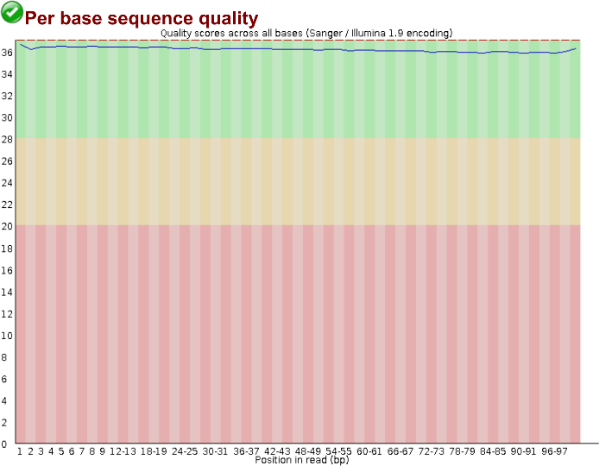

**Figure S1 (Cont.)**

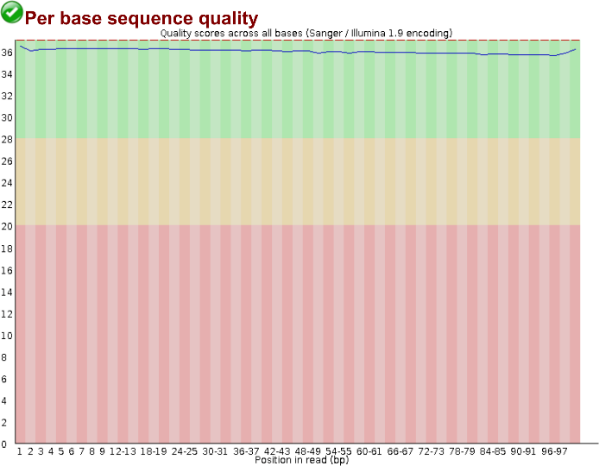

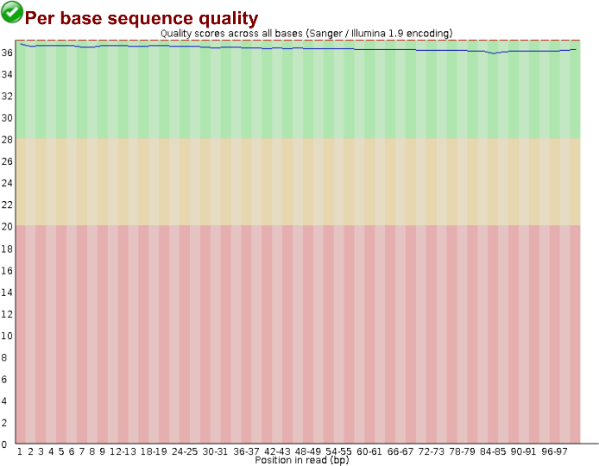
**D.	CD1-brain-M-4M-DNA-3**

**E.	CD1-brain-M-5M-DNA-5**

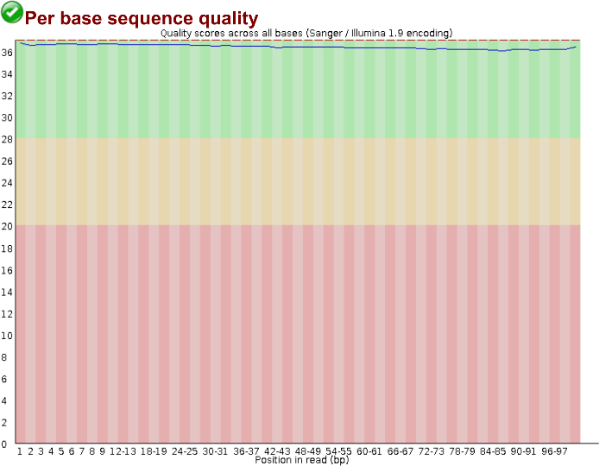

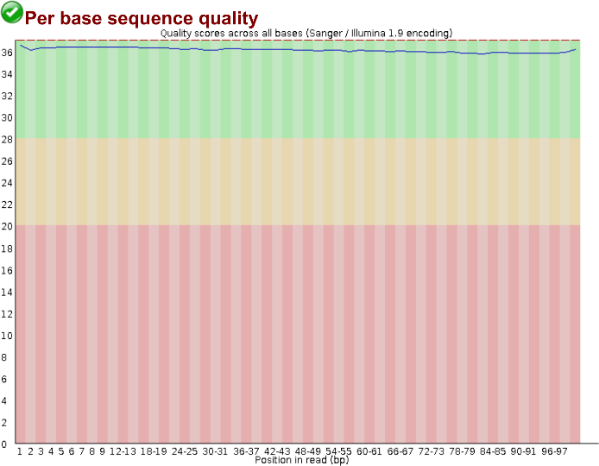

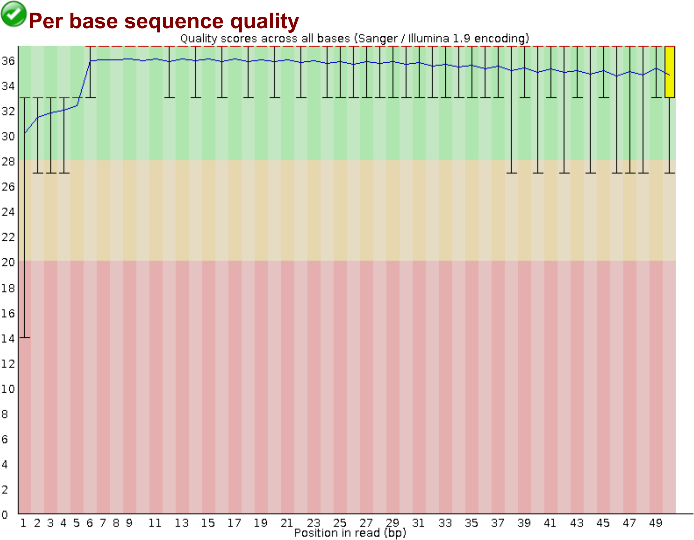
**F.	Cont-Adipose-ATACseq-Rep1**

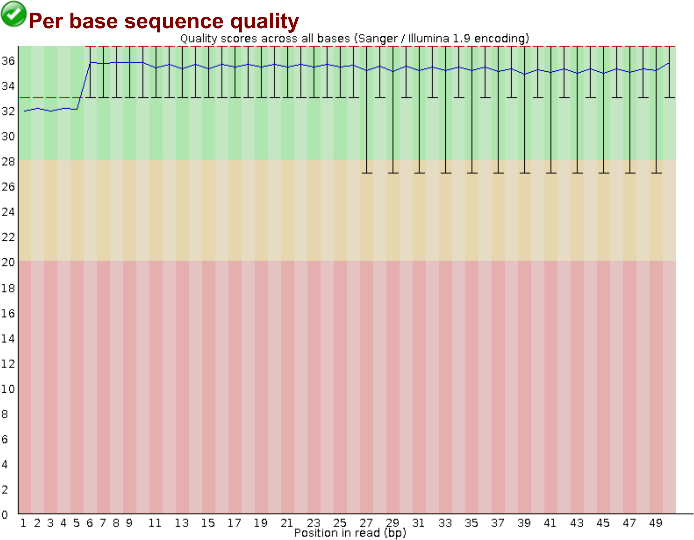

**Figure S1 (Cont.)**

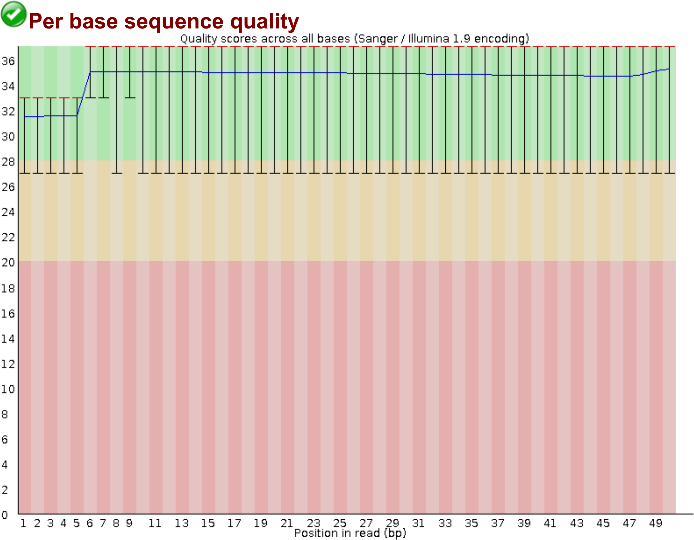

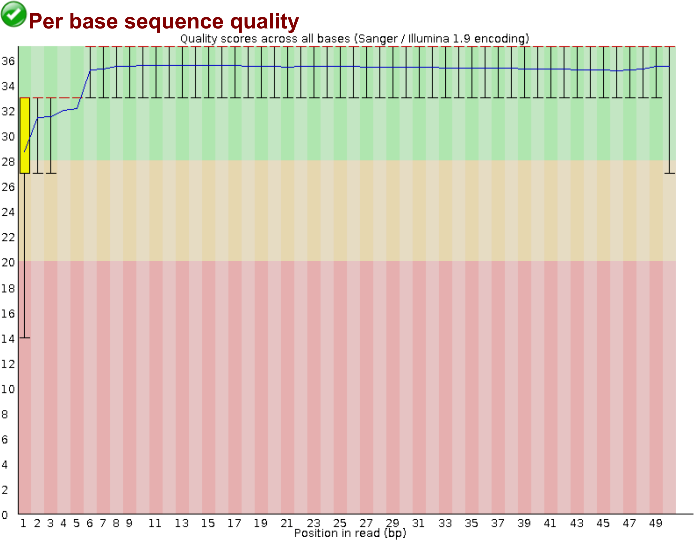
**G.	Cont-Adipose-ATACseq-Rep2**

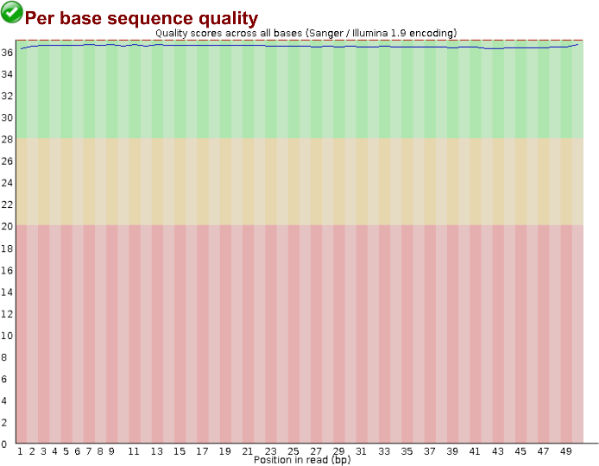

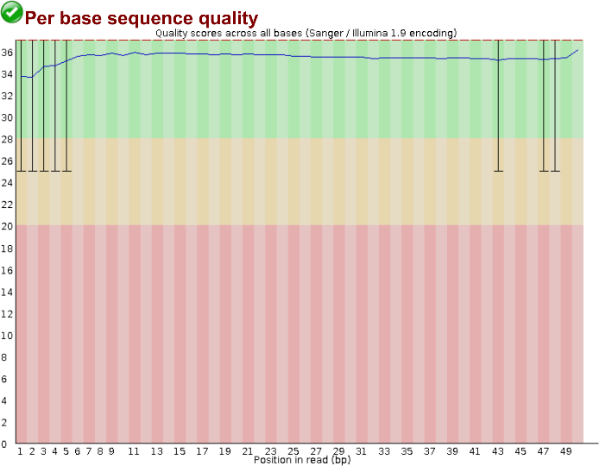
**H.	BPAF-Adipose-ATACseq-Rep1**

**I.	BPAF-Adipose-ATACseq-Rep2**

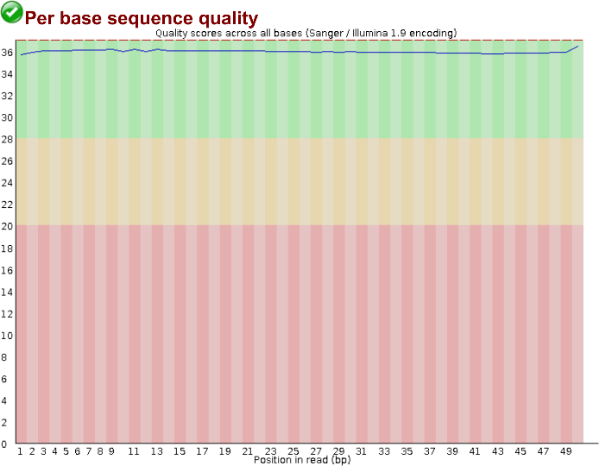

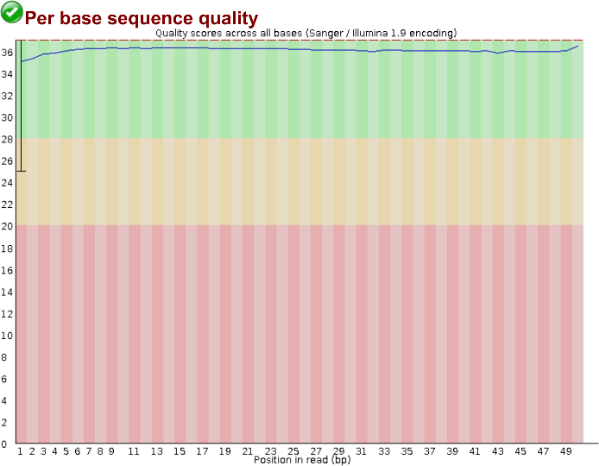

**Figure S1 (Cont.)**

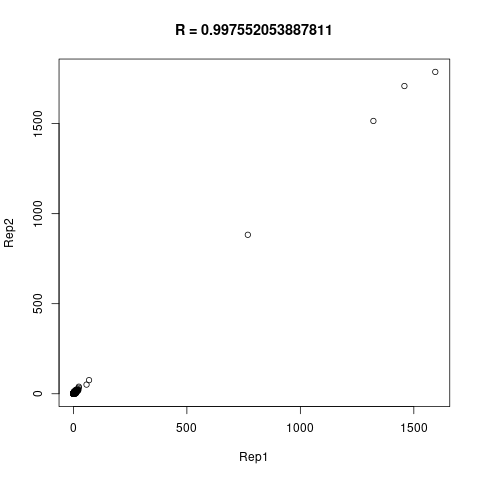

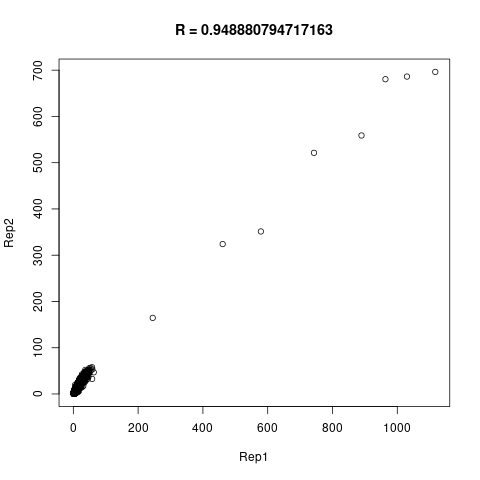
**J.** BPA	Control

**K.	Unfiltered Validation SNPs**

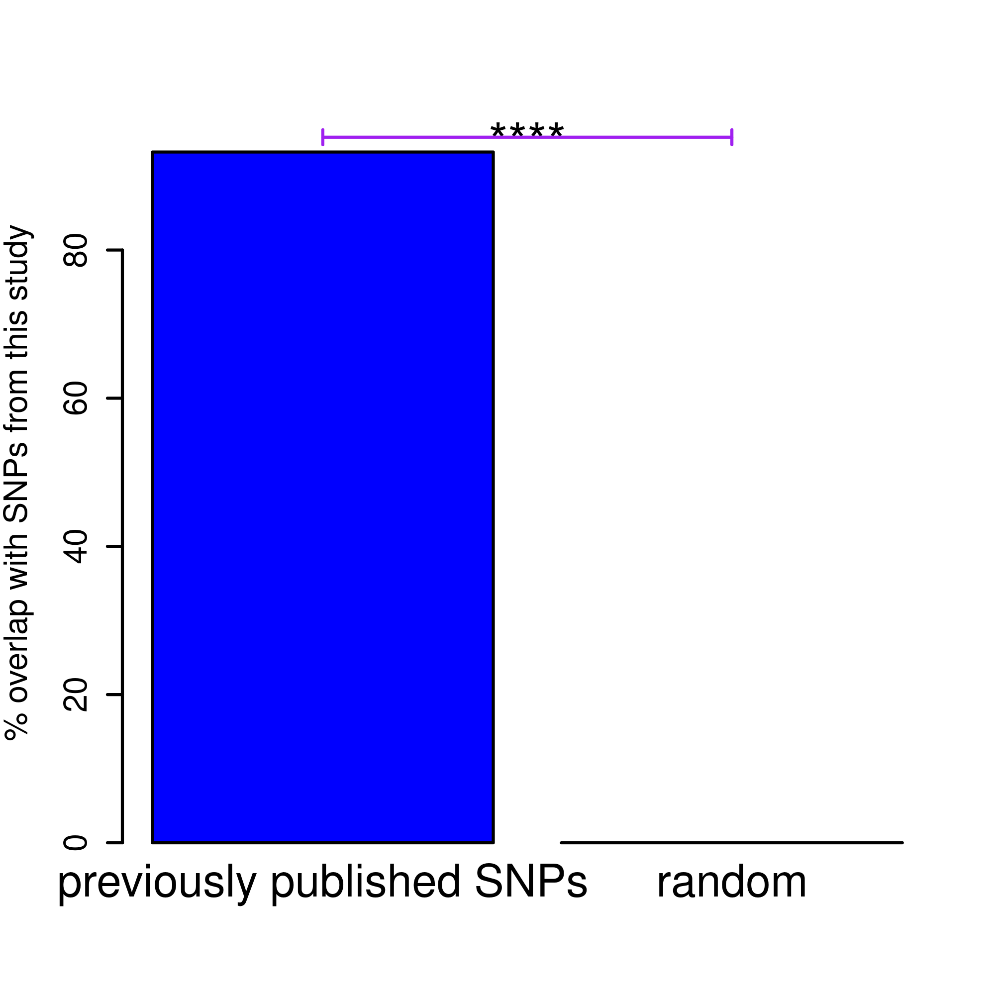

**Figure S1 (Cont.)**

**L.	Non-Uniform SNPs	Non-Uniform SNPs**

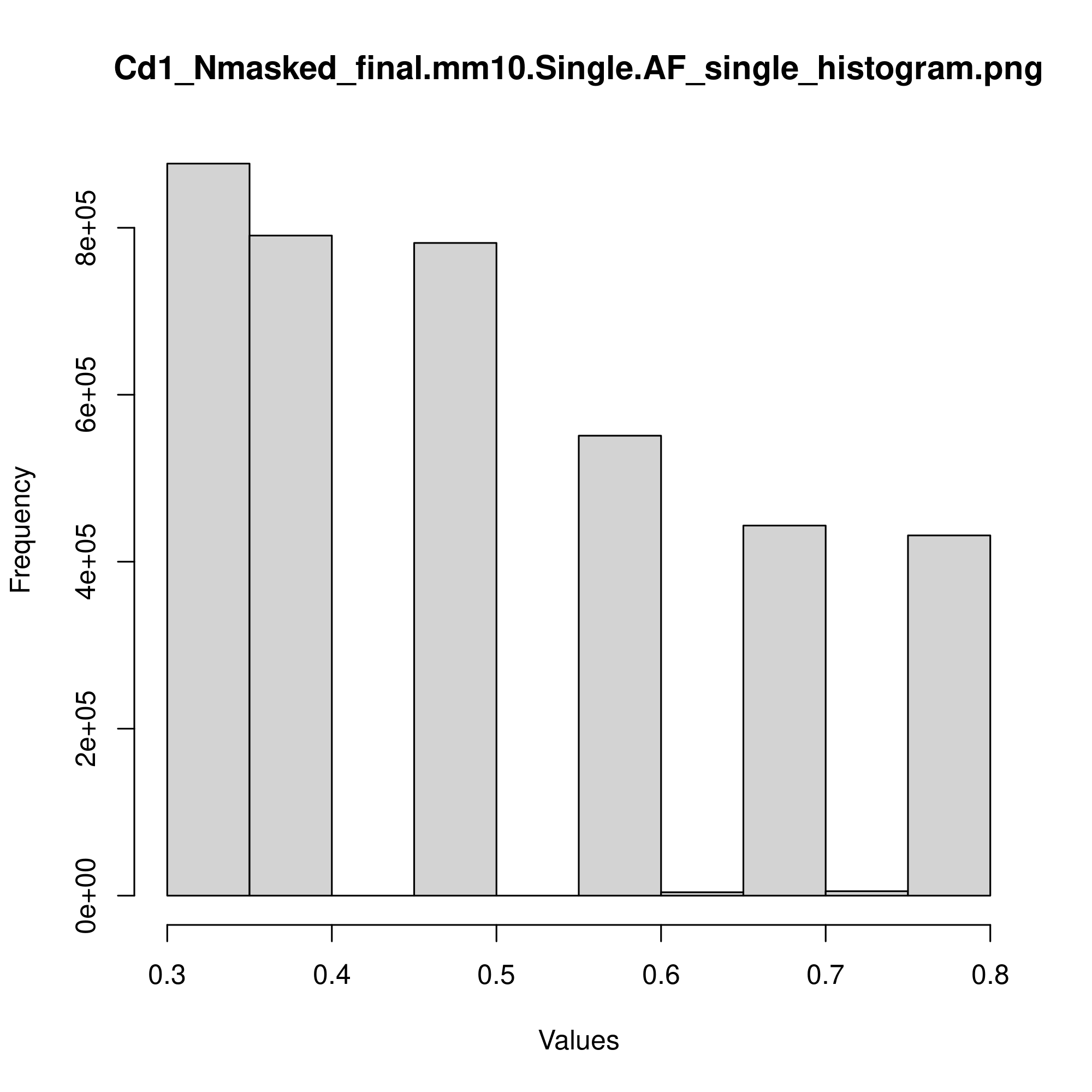

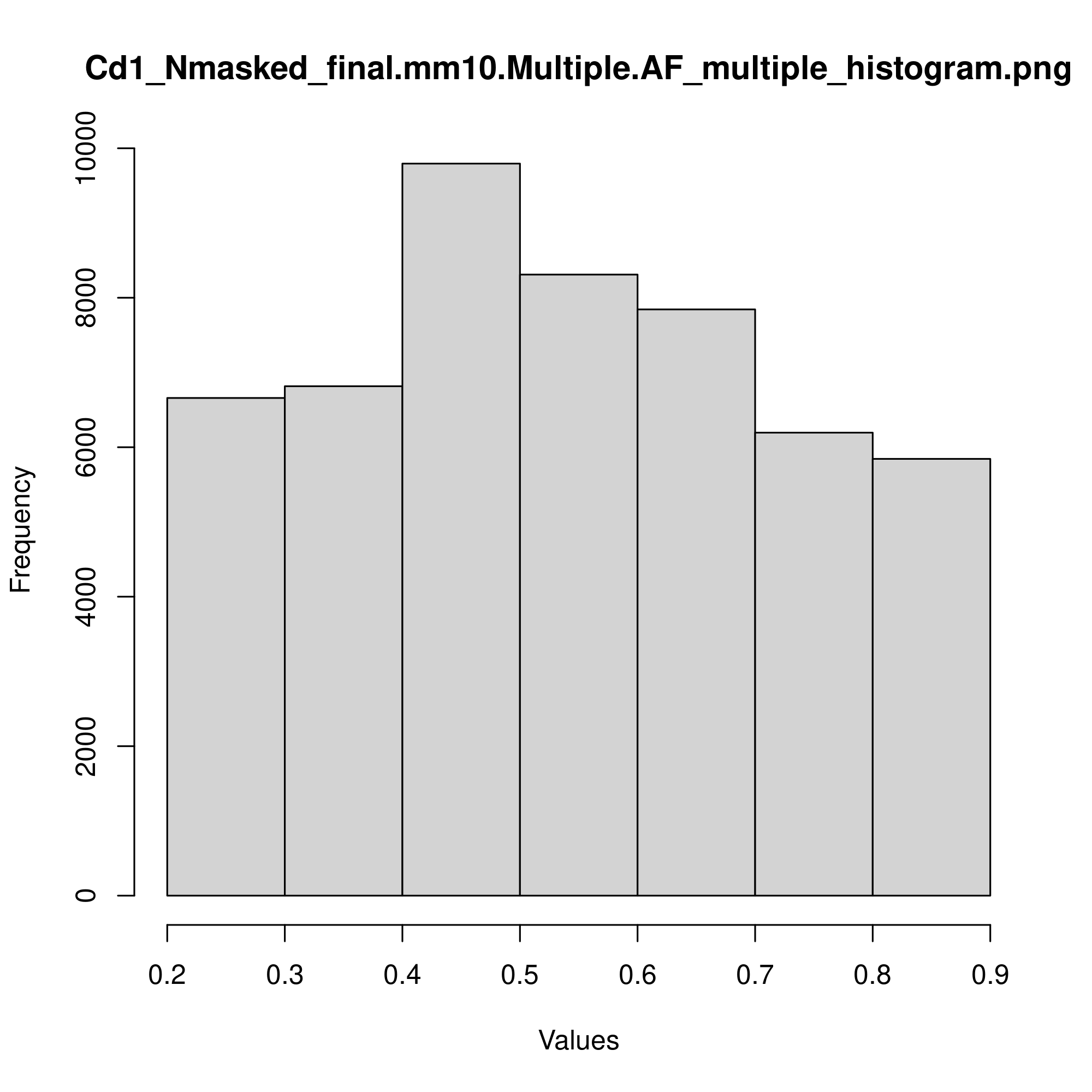
 **1 alt. allele	>1 alt. allele**

**Uniform Indels	Uniform SNPs**

**1 alt. allele	1 alt. allele**

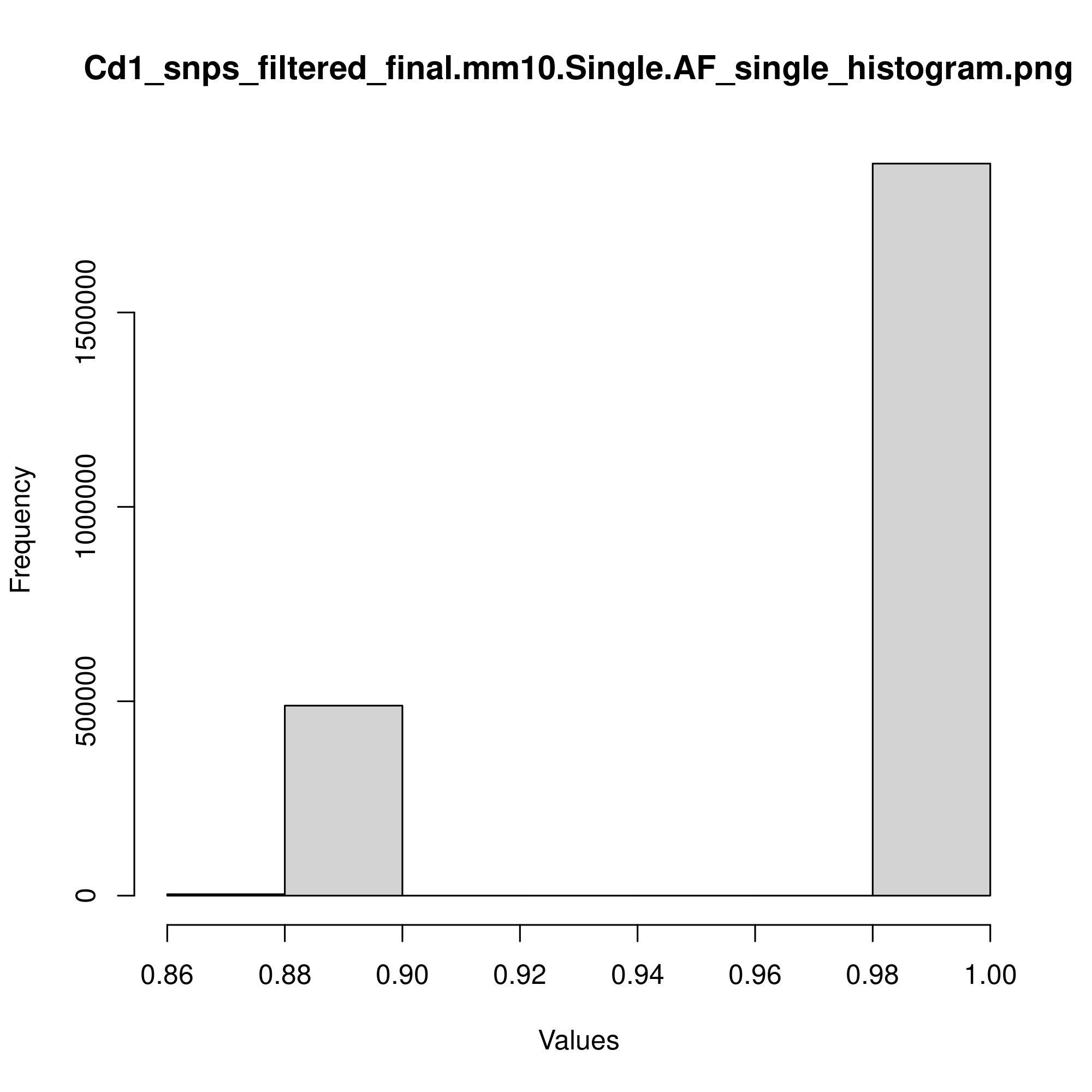

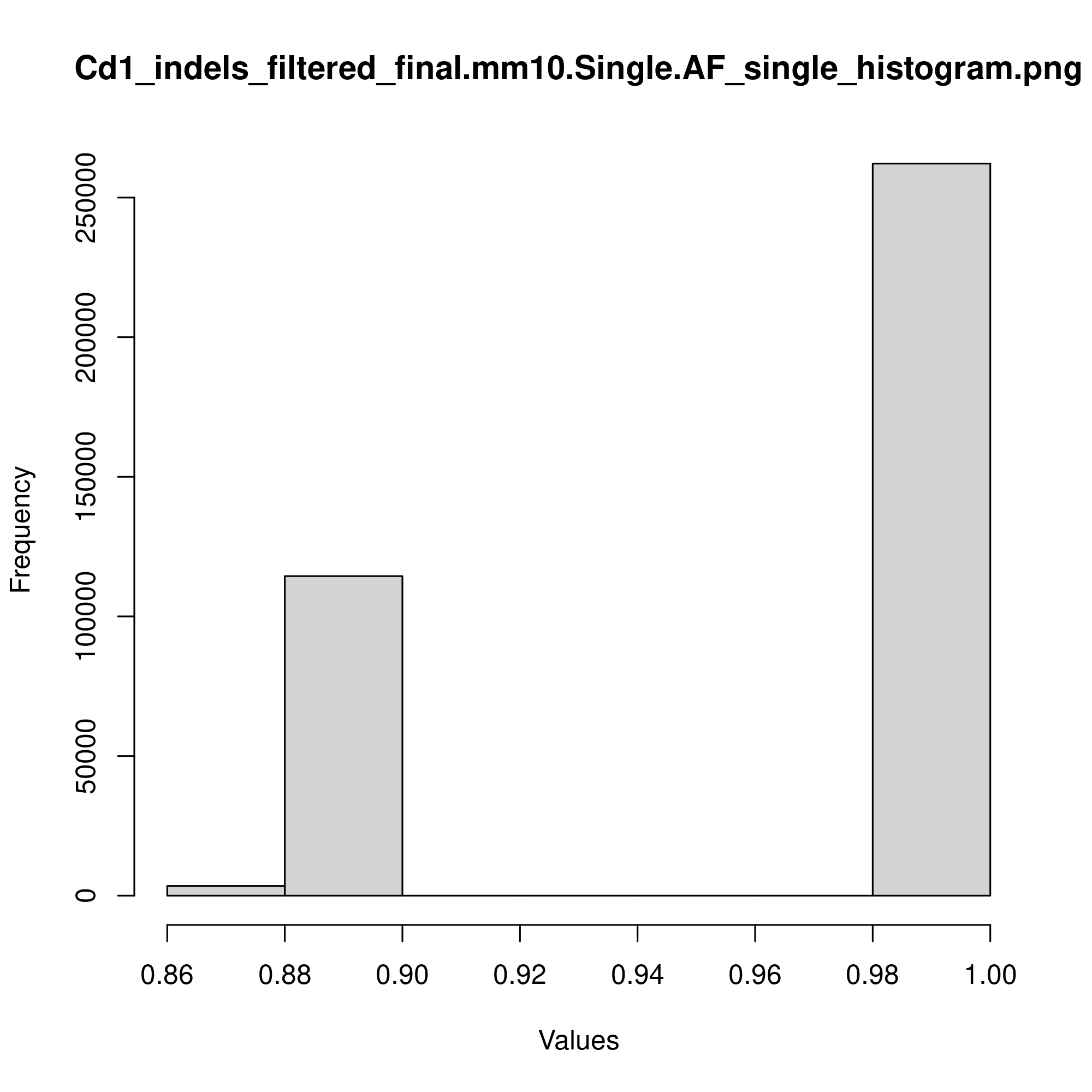

**Non-Uniform Indels	Non-Uniform Indels**

**1 alt. allele	>1 alt. allele**

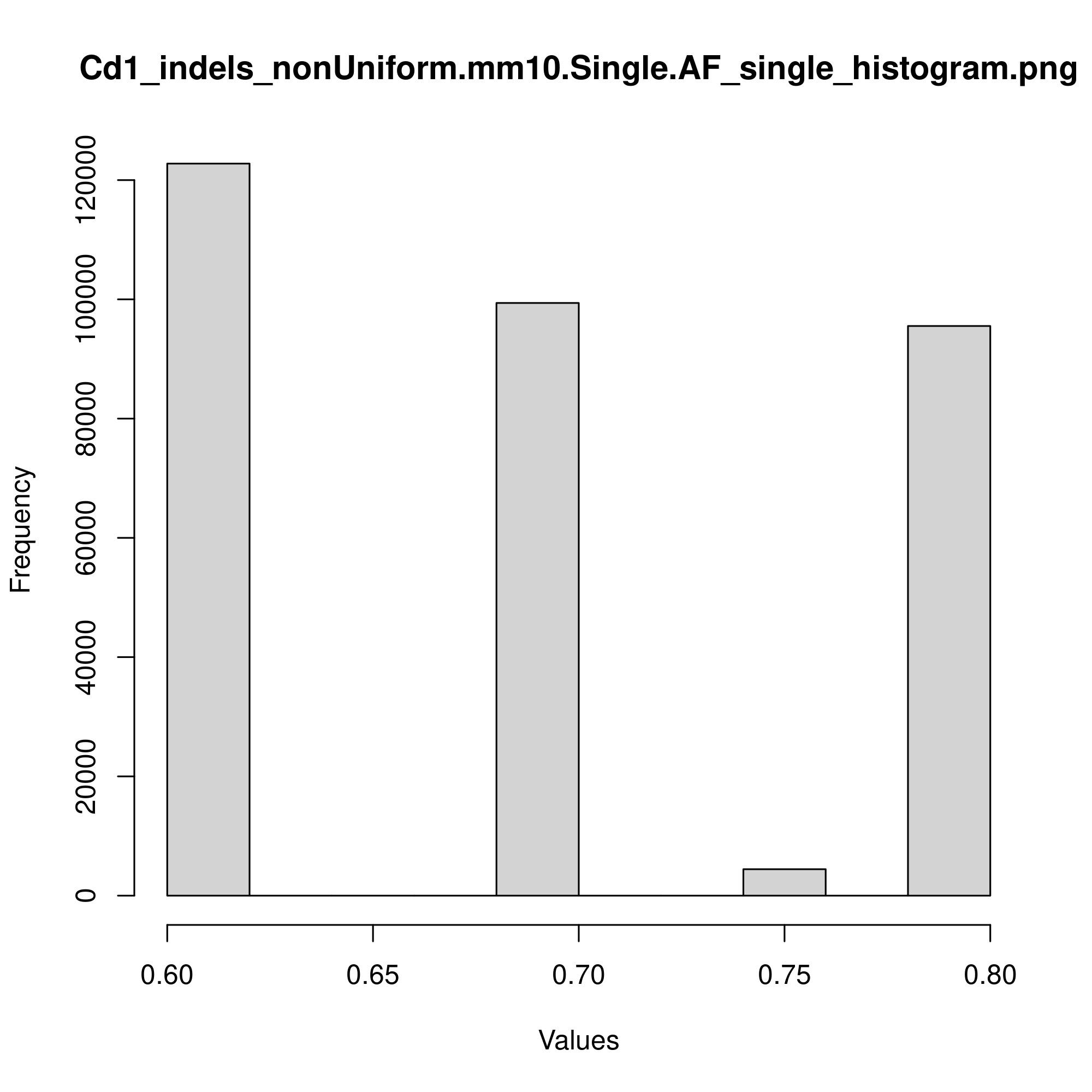

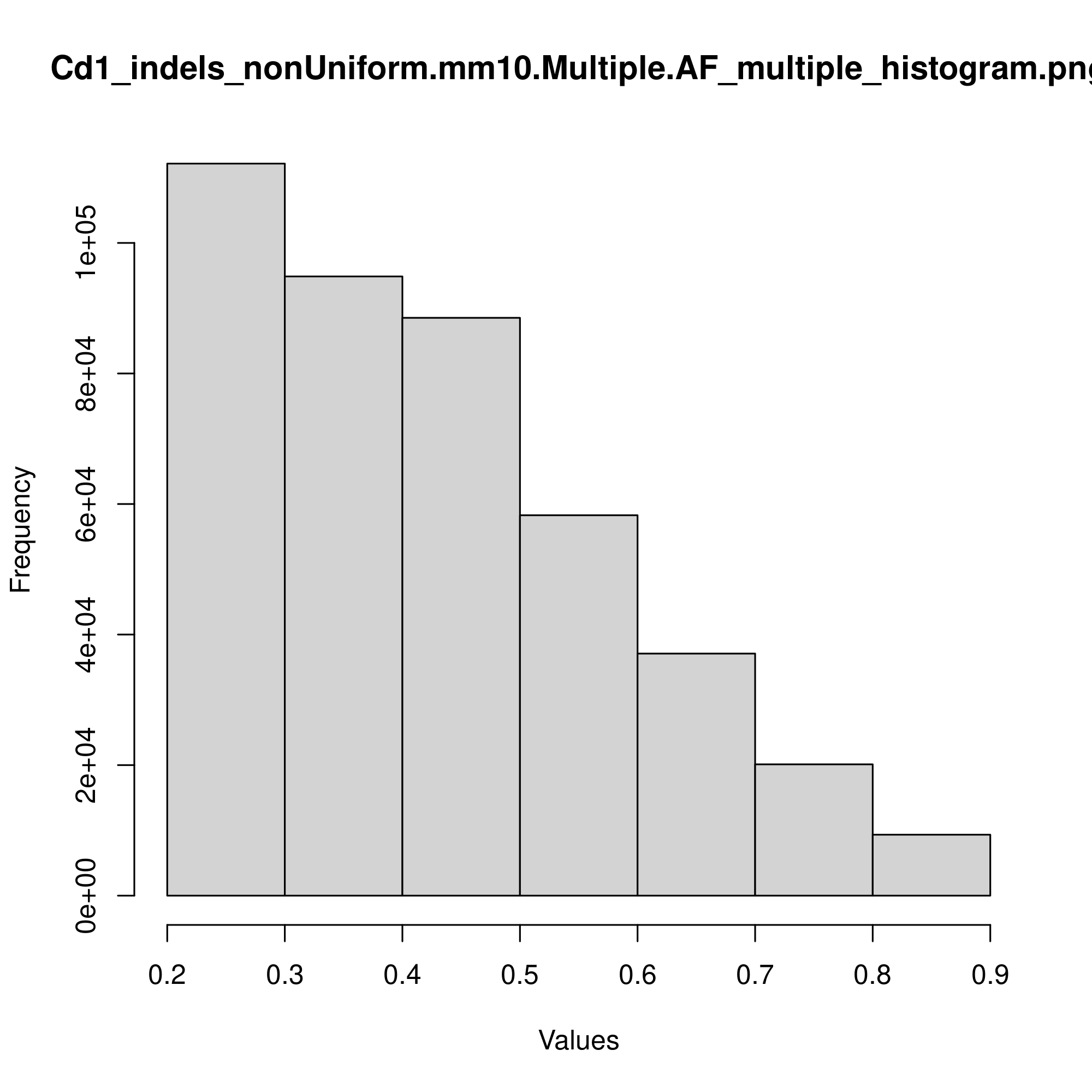

**Figure S1 (Cont.)**

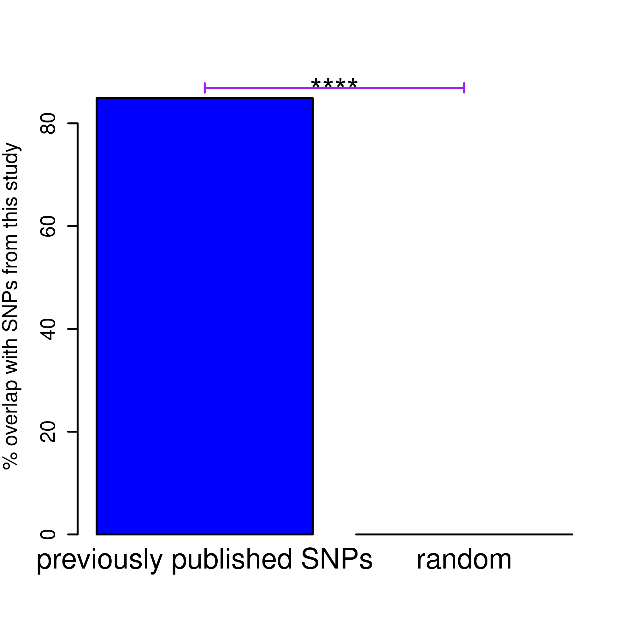

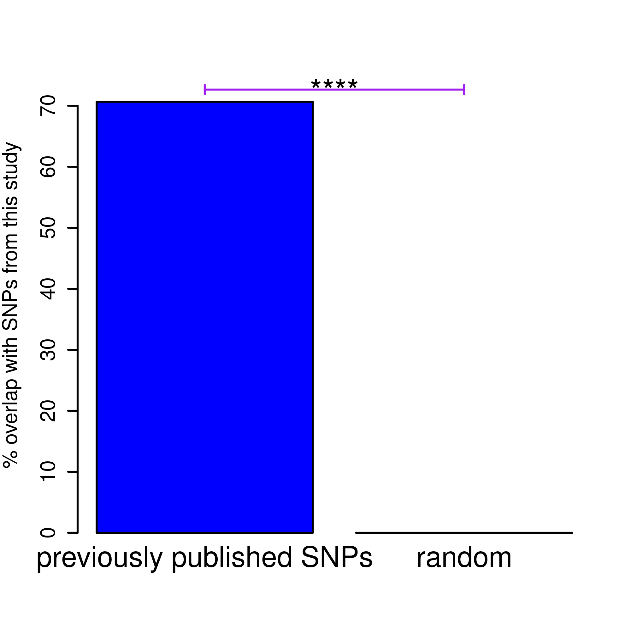
**M. Non-uniform	Uniform**

**Figure S1. Validation of sequencing data generated in this study. A-E** Plots showing the distribution of base quality scores at each position along the left and right read pair (left and right plots, respectively) for the indicated Cd1 gDNA sample. **F-I** Same as **A-E**, except for the indicated ATAC-seq sample. **J** Scatterplots comparing RPKM values of the two replicates of the indicated ATAC-seq sample generated in this study (BPA or Control). Pearson’s correlation coefficient is shown above each plot. **K** Barplot showing the percent overlap of the combined set of uniform and non-uniform, previously published SNPs that overlap with our SNPs before removal of low abundance and low frequency SNPs, and comparing that to the percent overlap of randomly shuffled regions with our SNPs before removal of low abundance and low frequency SNPs. **L** Histograms of frequencies of the most abundant alternative allele for the indicated variant type. 1 alt. allele = variants with only 1 alternative allele that differs from the reference (mm10) allele; >1 alt. allele = variants with more than one alternative allele that differs from the

**Figure S1 (Cont.)**

reference (mm10) allele. Categories not shown do not have any variants. **M** Same as K, except using just non-

uniform SNPs (left panel), and just uniform SNPs (right panel). P-values in this figure were calculated by Fisher’s exact test, with cutoffs shown as follows: * p < .01 ; ** p < .001 ; *** p < .00001 ; **** p < .0000000001.

**Figure S2**

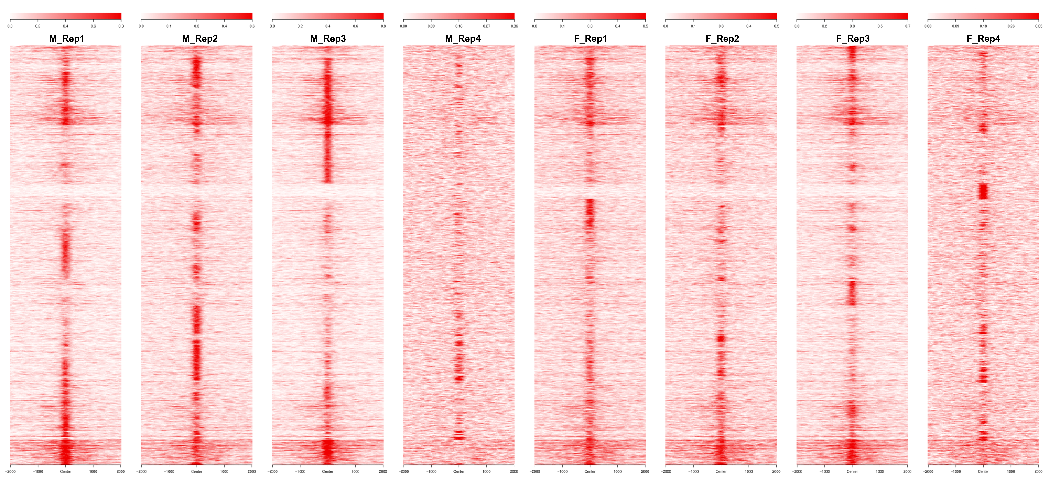
**A.**

N=9602

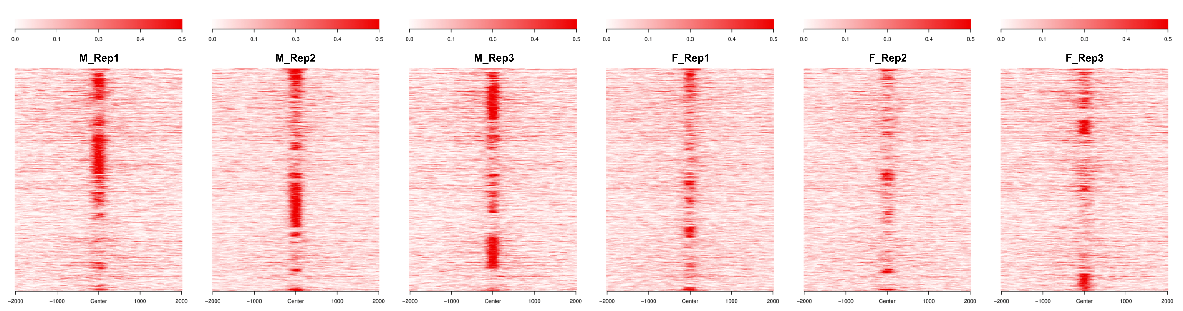
**B.**

N=3387

**Figure S2 (Cont.)**

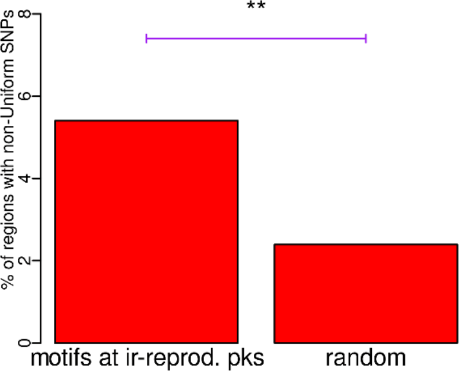

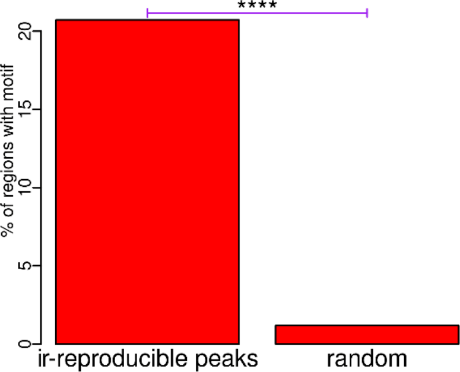
**C.	D.**

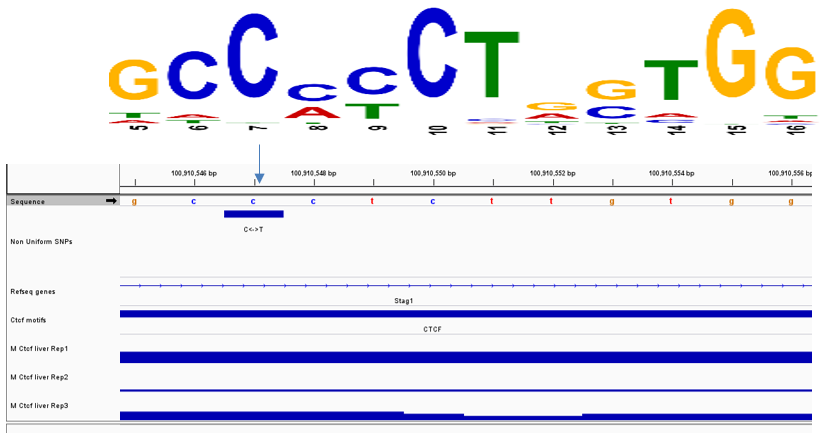
**E.**

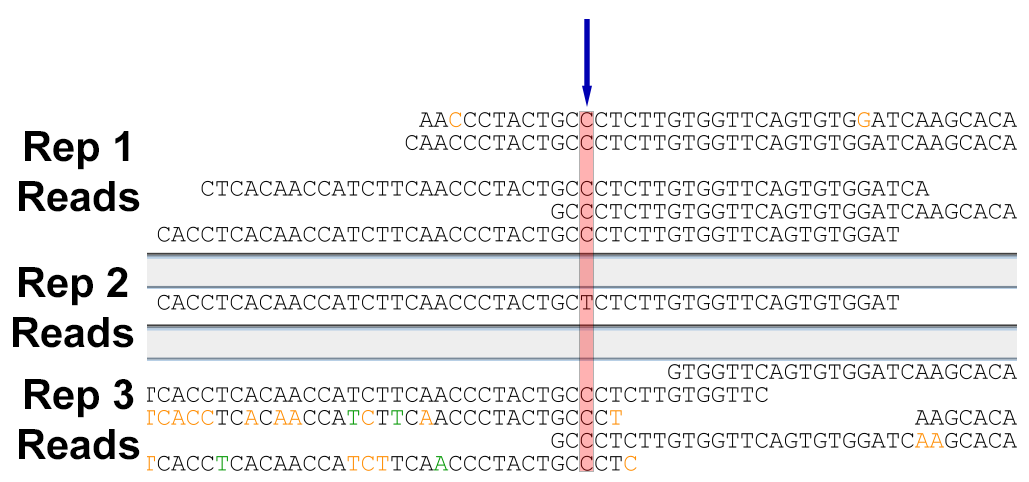
**F.**

**Figure S2 (Cont.)**

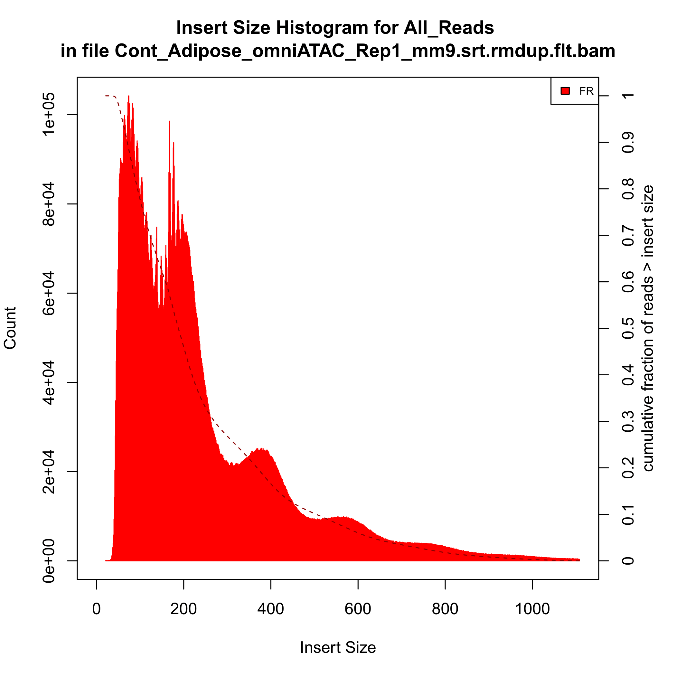

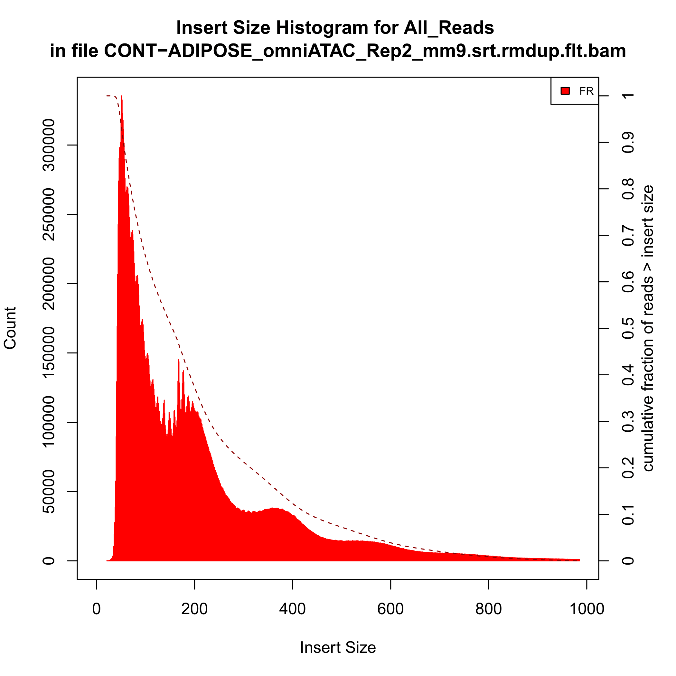
**G.**

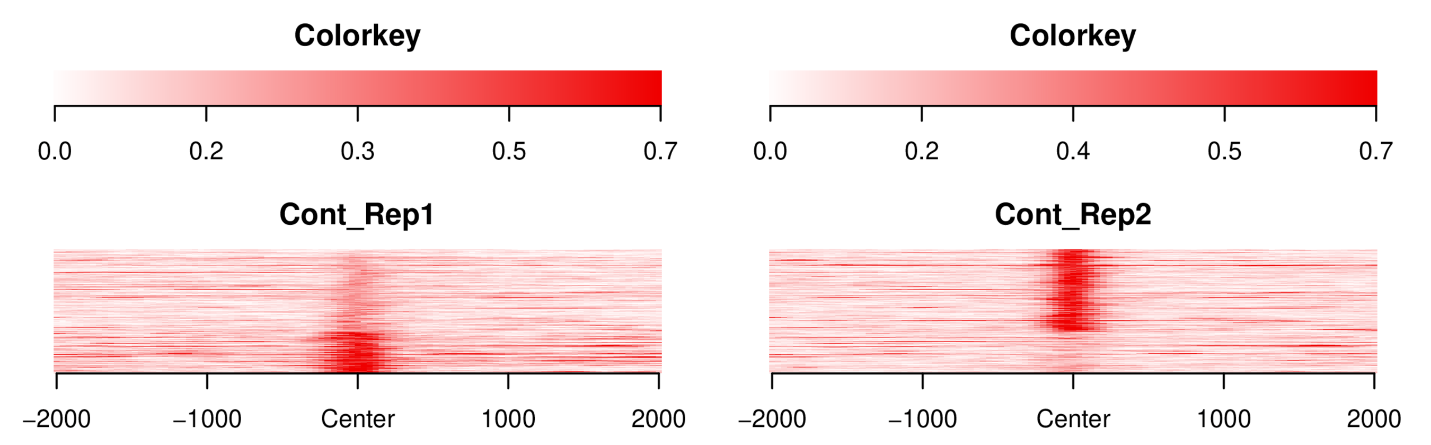
**H.**

N=521

**Figure S2. Related to Figure 1. A** Heatmaps showing the enrichment of Ctcf ChIP-seq signal at ir-reproducible peaks when all four replicates from both male and female liver samples were considered. **B** Same as **A,** except in this case, only considering replicates 1-3 of both the male and female samples. **C** Barplots comparing the percentage of ir-reproducible peaks (with replicates 1-3 of both male and female samples considered) that contain a called Ctcf motif to the ratio of randomly shuffled

**Figure S2 (Cont.)**

regions that contain a Ctcf motif. **D** Barplots comparing the percentage of ir-reproducible peaks (with replicates 1-3 of both male and female samples considered) with a Ctcf motif that overlap with a non-uniform SNP to the percentage of randomly shuffled regions that overlap a non-uniform SNP. **E** Genome browser image of Reads per million (RPM)-normalized Ctcf ChIP-seq coverage around a non-uniform SNP that occurs at a highly conserved C in a called Ctcf motif (reverse complimented). The reverse-compliment CTCF motif is pictured at top, and the arrow points to the nucleotide with the non-uniform SNP. **F** The sequenced Ctcf ChIP-seq reads from the indicated replicate that map to the non-uniform SNP displayed in Figure S1E. Nucleotides are colored according to their phred sequencing quality score as follows: black for 30 or above, orange < 30, green < 20, blue < 10. **G** Histograms of the fragment length distribution for the two replicates of adipose ATAC-seq in control, CD-1 mice. **H** Heatmaps showing enrichment of ATAC-seq signal at ir-reproducible peaks. P-values in this figure were calculated by Fisher’s exact test, with cutoffs shown as follows: * p < .01 ; ** p < .001 ; *** p < .00001 ; **** p < .0000000001.

**Figure S3**

**A.**
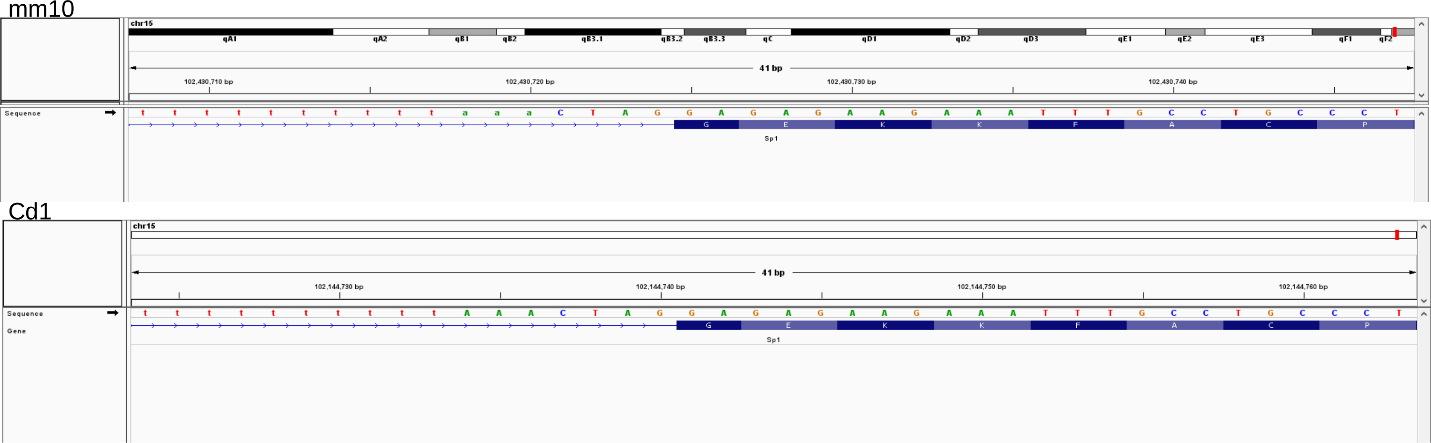

**B.**
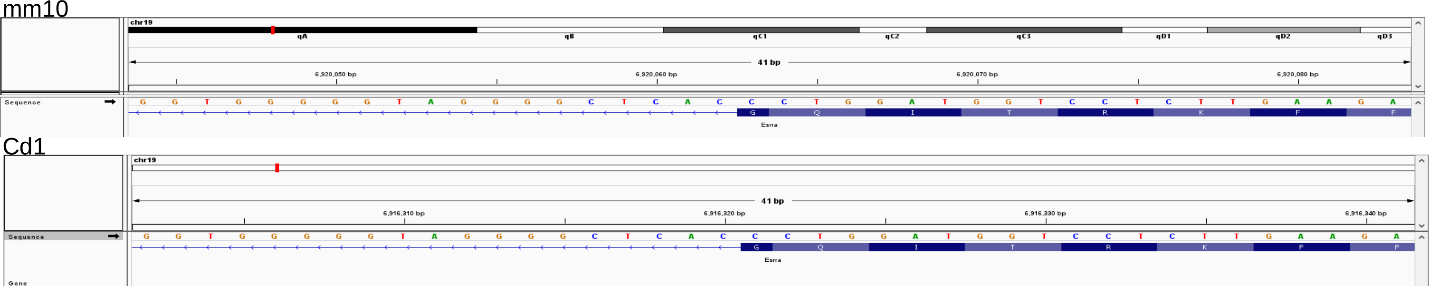

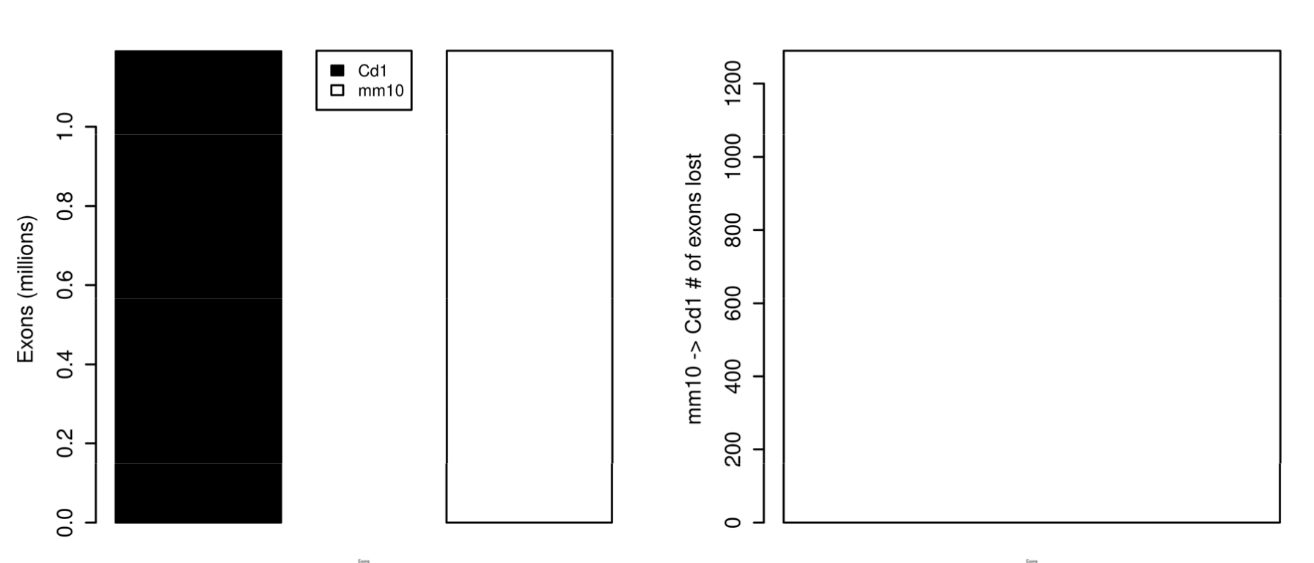
**C.**

**Figure S3 (Cont.)**

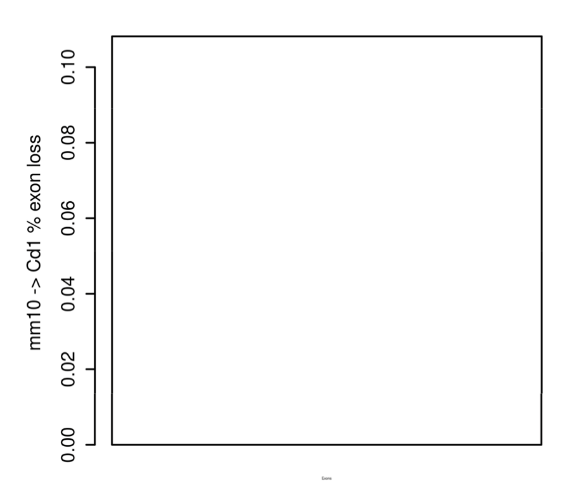
**D.**

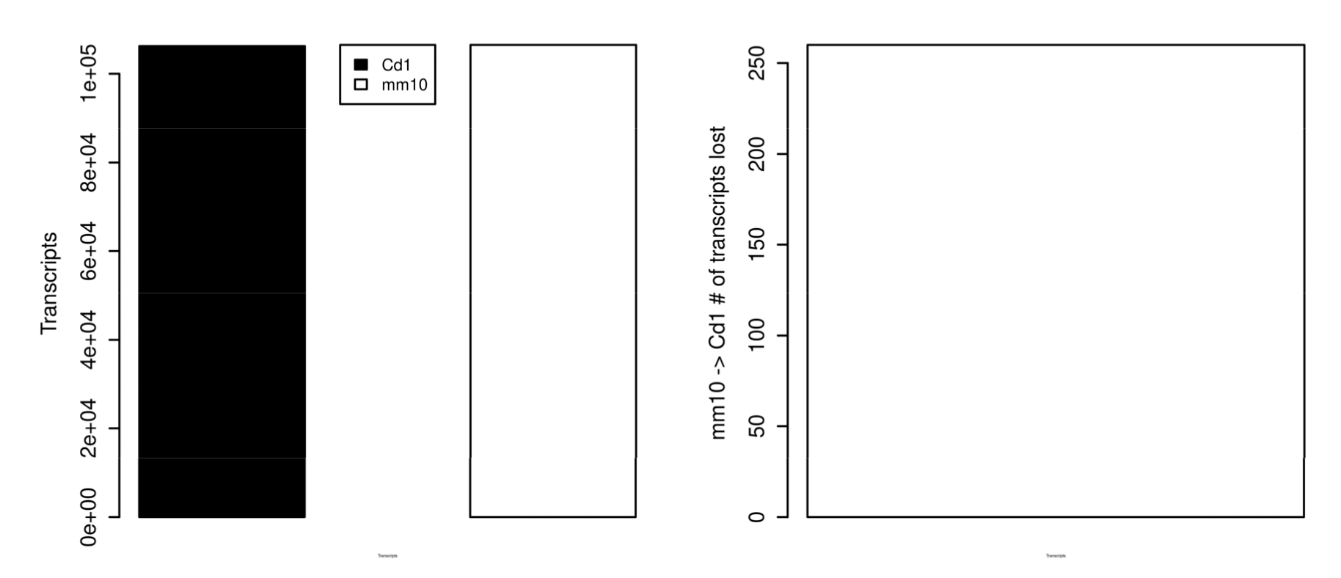
**E.**

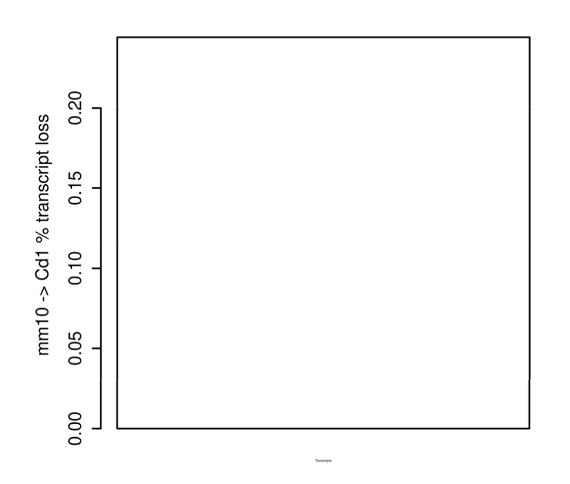
**F.**

**Figure S3 (Cont.)**

**Figure S3. Validation of the newly generated CD-1 gene annotations. A** Genome browser views in mm10 coordinates (top) and Cd1 coordinates (bottom) around an intron-exon junction of the Sp1 gene. Both the mm10 and Cd1 nucleotide sequences at this region are displayed for comparison. **B** Same as **A**, except for an intron-exon junction of the Esrra gene.

**Figure S4**

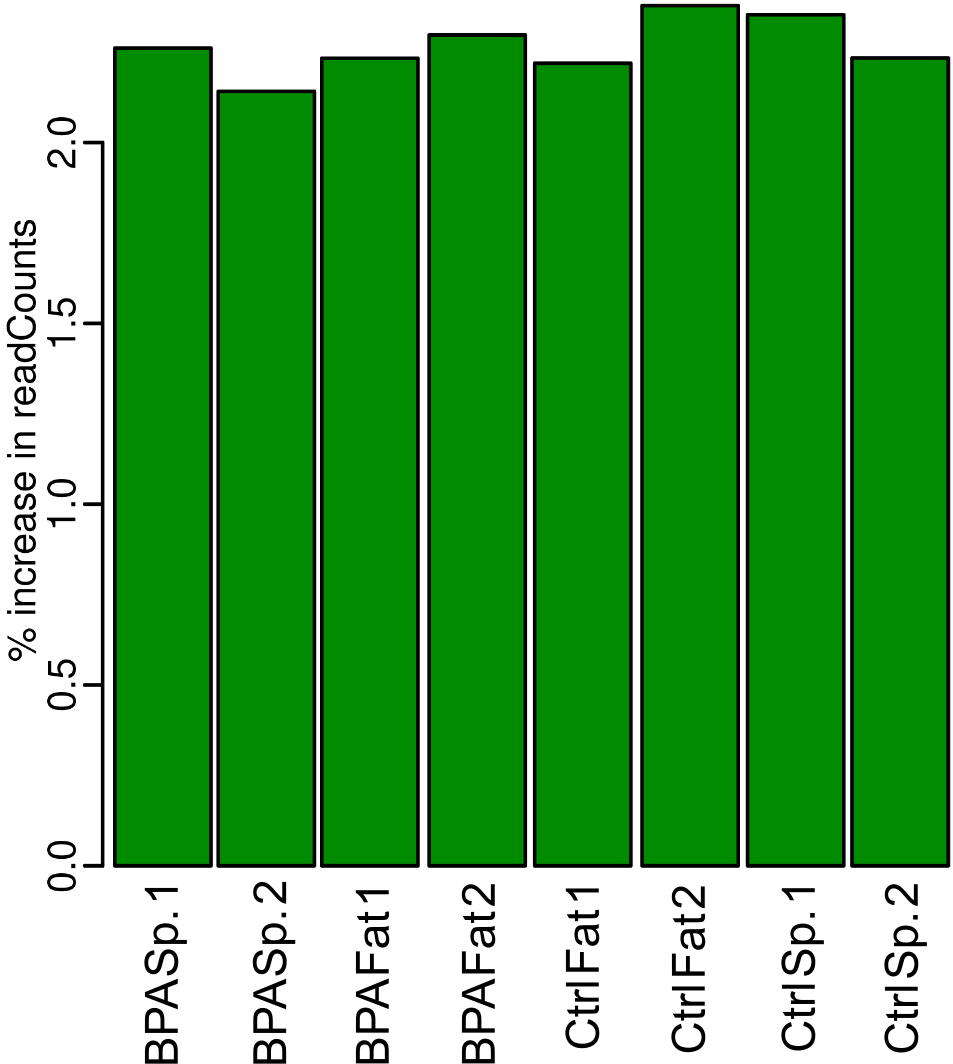
**A.**

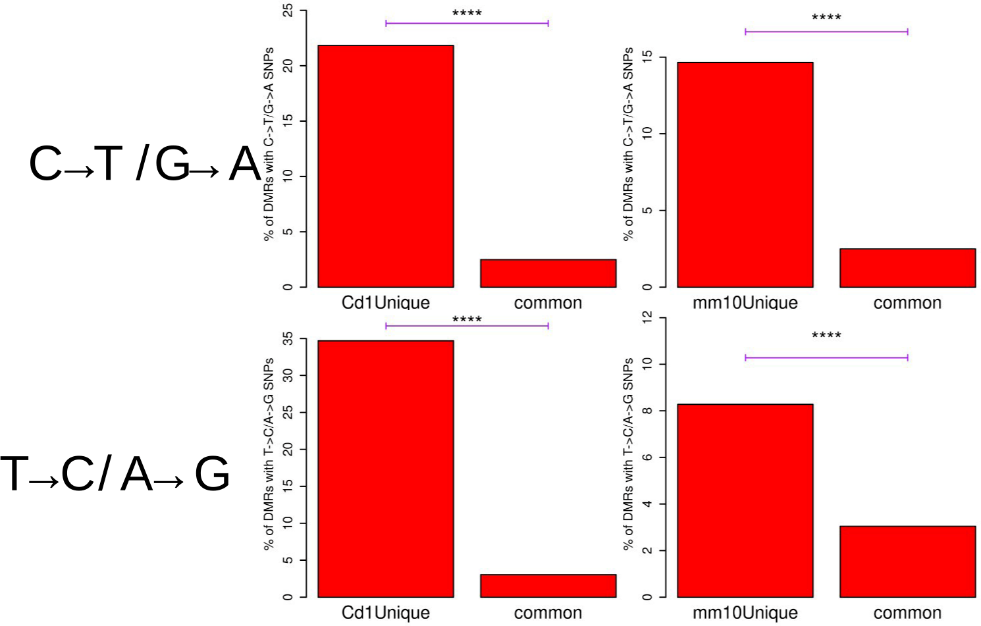
**B.**

**Figure S4. Related to Figure 3. A** Barplots showing the percent increase in the number of nonduplicate BS-seq reads mapped when mapping to Cd1 rather than mm10, for the indicated

**Figure S4 (Cont.)**

samples. Positive values indicate more reads mapped in Cd1 than mm10. **B** Barplots comparing the percentage of the indicated DMR sets overlapping with a uniform C->T or G->A SNP (mm10->Cd1; top row), or comparing the indicate DMR sets overlapping with a uniform T->C or A->G SNP (mm10->Cd1; bottom row). Cd1Unique: DMRs that are called when mapping to Cd1 but not when mapping to mm10; mm10Unique: DMRs that are called when mapping to mm10 but not when mapping to Cd1; common: DMRs that are called both when mapping to Cd1 and to mm10. P-values in this figure were calculated by Fisher’s exact test, with cutoffs shown as follows: * p < .01 ; ** p < .001 ; *** p < .00001 ; **** p < .0000000001.

**Figure S5**

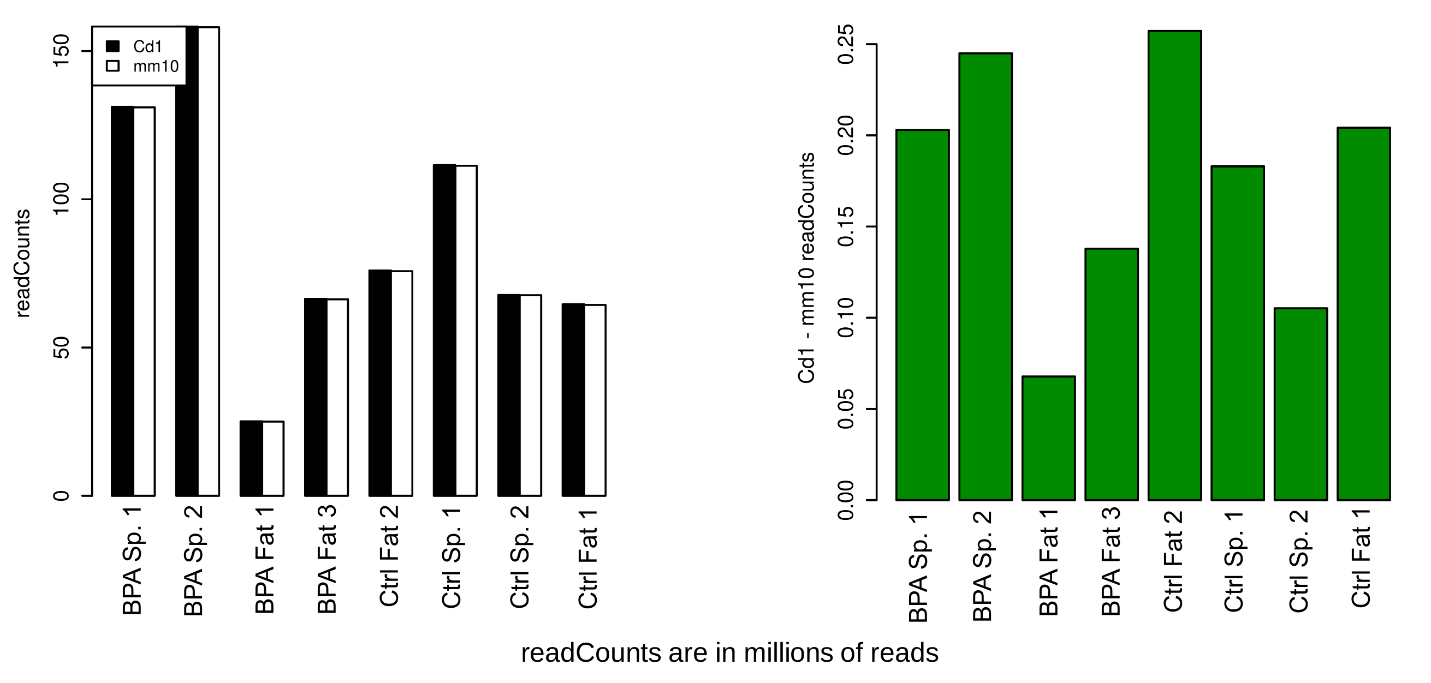
**A.**

**B.**

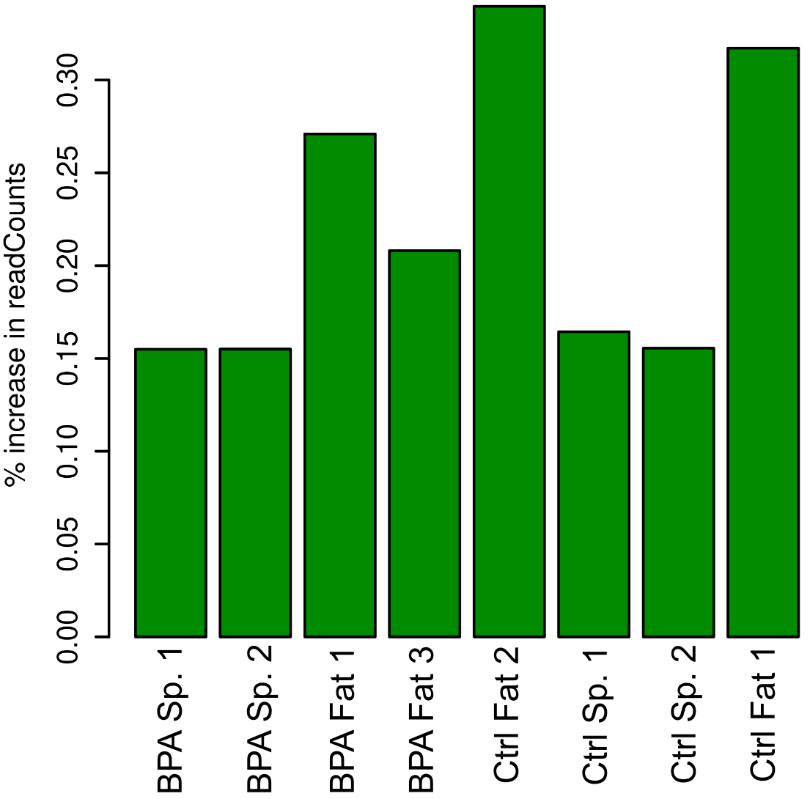

**Figure S5. Related to Figure 4. A** Left panel: Barplots showing the number of non-duplicate ATAC-seq reads mapped to either Cd1 or mm10 from the indicated samples. Right panel: Barplots

**Figure S5 (Cont.)**

showing the difference between the number of nonduplicate ATAC-seq reads mapped in Cd1 and mm10. Positive values indicate more reads mapped in Cd1 than mm10. **B** Barplots showing the percent increase in the number of nonduplicate ATAC-seq reads mapped when mapping to Cd1 rather than mm10. Positive values indicate more reads mapped in Cd1 than mm10.
